# Biomaterial Scaffold Stiffness Influences the Foreign Body Reaction, Tissue Stiffness, Angiogenesis and Neuroregeneration in Spinal Cord Injury

**DOI:** 10.1101/2024.04.02.587745

**Authors:** Yifeng Zheng, Maximilian Nützl, Thomas Schackel, Jing Chen, Norbert Weidner, Rainer Müller, Radhika Puttagunta

## Abstract

Biomaterial scaffold engineering presents great potential in promoting axonal regrowth after spinal cord injury (SCI), yet persistent challenges remain, including the surrounding host foreign body reaction and improper host-implant integration. Recent advances in mechanobiology spark interest in optimizing the mechanical properties of biomaterial scaffolds to alleviate the foreign body reaction and facilitate seamless integration. The impact of scaffold stiffness on injured spinal cords has not been thoroughly investigated. Herein, we introduce stiffness-varied alginate anisotropic capillary hydrogel scaffolds implanted into adult rat C5 spinal cords post-lateral hemisection. Four weeks post-implantation, scaffolds with a stiffness approaching that of the spinal cord effectively minimize the host foreign body reaction via yes-associated protein (YAP) nuclear translocation. Concurrently, the softest scaffolds maximize cell infiltration and angiogenesis, fostering significant axonal regrowth but limiting the rostral-caudal linear growth. Furthermore, as measured by atomic force microscopy (AFM), the surrounding spinal cord softens when in contact with the stiffest scaffold while maintaining a natural level in contact with the softest one. In conclusion, our findings underscore the pivotal role of stiffness in scaffold engineering for SCI *in vivo*, paving the way for the optimal development of efficacious biomaterial scaffolds for tissue engineering in the central nervous system.

## 1. Introduction

Spinal cord injury (SCI) frequently leads to the disruption of the unique and complex bridge connecting the brain and periphery, ultimately resulting in various neurological deficits.[1] Following SCI, the growth ability of axons is limited but can be enhanced through regenerative strategies.[2] Regrowing axons face significant challenges in traversing the lesion and establishing reinnervation with distal targets, extensively due to the inhibitory lesion environment.[3] Biomaterial scaffolds exhibit great promise in bridging the lesion by integrating into the spinal cord physically. Previously, we reported an anisotropic capillary hydrogel (ACH) scaffold comprised of the heteromeric polysaccharide alginate, facilitating linear axonal regrowth upon *in vivo* implantation.[4] Furthermore, the scaffold capillaries can be infused with extracellular matrix (ECM) proteins and/or functional cells to enhance the growth environment.[5] Nevertheless, being a foreign implant, the scaffold encounters natural resistance from the host spinal cord, triggering a foreign body reaction at the host-implant interface, characterized by inflammation and fibrosis.[6] It generally separates the implanted scaffold from the host tissue and strongly limits the host-implant integration as well as further axonal regrowth.

Recent advances in mechanobiology have sparked interest in exploring the role of mechanics in various pathological disease processes.[7] *In vitro*, alterations in the stiffness of the environment dictates the behavior and function of various cell types, including glial cells,[8] neurons,[9] macrophages,[10] fibroblasts,[11] and stem cells.[12] When cultured on a stiffer substrate, glial cells typically undergo gliosis, characterized by morphological changes and upregulation of inflammatory genes and proteins.[8, 13] Furthermore, macrophages transition towards a pro-inflammatory phenotype with impaired phagocytosis when exposed to a stiff matrix compared to a soft one.[10, 14] Additionally, various neuronal cells and their axons exhibit mechanosensitivity, responding to changes in substrate stiffness both *in vitro* and *in vivo*.[9, 15] Mechanotransduction, the process of translating mechanical cues into biological signals, is believed to engage intricate mechanisms, including the F-actin and RhoA signaling pathways.[16] In mechanotransduction pathways, the yes-associated protein (YAP) is recognized as a key downstream mediator in various cell types,[11, 14a, 17] however its role in complex tissue interactions remains unexplored.

Tissue stiffness plays a crucial role in embryonic development, sprouting angiogenesis, and potentially neural repair.[17b, 18] During the measurement of tissue stiffness, variations in dimension scales result in significant heterogeneity. Micro-indentation studies of the spinal cord reported elastic Young’s modulus less than 1 kPa,[19] whereas macro-tensile studies reported three orders of magnitude higher (1.52 MPa).[20] However, while bulk stiffness ensures *in vivo* stability, microscale properties are more essential in regards to regulating cellular behavior and further biological events.[21] After injury, the spinal cord undergoes softening, associated with an enhanced inflammatory reaction and ECM protein deposition.[19b] Nevertheless, the tissue stiffness of the spinal cord when in contact with a biomaterial scaffold has never been investigated. In recent times, atomic force microscopy (AFM) has emerged as a tool for precise mechanical characterization of *ex vivo* living tissues at microscale and/or nanoscale.[22] Thus allowing measurement of host-implant interface mechanics following implantation.

While support for the role of mechanical stiffness in regulating the foreign body reaction and neuronal growth continues to build,[23] it is unclear if the stiffness of biomaterial scaffolds have significant implications for the efficacy of tissue engineering in the central nervous system (CNS). For example, it has been reported that the foreign body reaction was alleviated upon contact with a softer compliant polyacrylamide hydrogel implant in the brain.[8] However, in the more pliable spinal cord, with similar stiffness to the brain, it remains to be seen how soft the scaffold must be to limit foreign body reaction and yet retain its structure integrity for bridging the lesion site and physically guiding long distance axonal growth. In this study, we introduce stiffness-varied alginate hydrogel scaffolds implanted into adult rat C5 spinal cords post-lateral hemisection. Four weeks post-implantation, we examined the foreign body reaction and related mechanisms to scaffold stiffness, as well as host-implant integration through cell infiltration, angiogenesis, axonal regrowth and directionality within the biomaterial. Lastly, we questioned whether the host spinal tissue changes its mechanical properties (via AFM) in response to implant stiffness.

## 2. Results and Discussion

To investigate the significance of biomaterial scaffold stiffness for the foreign body reaction and axonal growth capacity in SCI, we developed stiffness-varied scaffolds approaching that of the spinal cord using our previously reported medical-grade alginate ACH.[4–5, 24] Given the architecture of the spinal cord white matter with long descending motor and ascending sensory tracts that are disrupted following injury, which require long-distance regeneration to re-establish target innervation for functional recovery, we have established the use of physical guidance cues through anisotropic capillary hydrogels for this purpose. Structurally, the oriented capillary guidance is created by the directional diffusion and complexation of divalent cations (Ca^2+^) within an aqueous sodium alginate solution (**Figure 1A**). Subsequently, chemical crosslinking with hexamethylene-diisocyanate (HDI) establishes covalent junction zones inside the physical hydrogel structure, strengthening the chemical and mechanical stability of the capillaries. To alter the mechanical properties of the ACH, we varied the alginate (0.5-1%) and HDI concentrations (5-50 mM), resulting in 4 distinct types, referred to as type I - IV (Figure 1B). In rheological measurements, the storage modulus (*G’*) of these ACH blocks increased from 0.4 kPa for type I gradually to 17 kPa for type IV (Figure 1C). This stiffness variation was further examined in 2×2×2 mm cuboids by AFM, which microscopically indented the surface parallel to the capillaries that would be directly in contact with the host tissue after implantation. The measured force-distance curves were fitted with a Hertz model to calculate the apparent Young’s modulus (*E*), providing a measure of elastic stiffness (Figure S1, Supporting Information). The quantified apparent Young’s modulus from type I to IV ranged from 1 to 9 kPa (Figure 1D-E).

**Figure 1:**
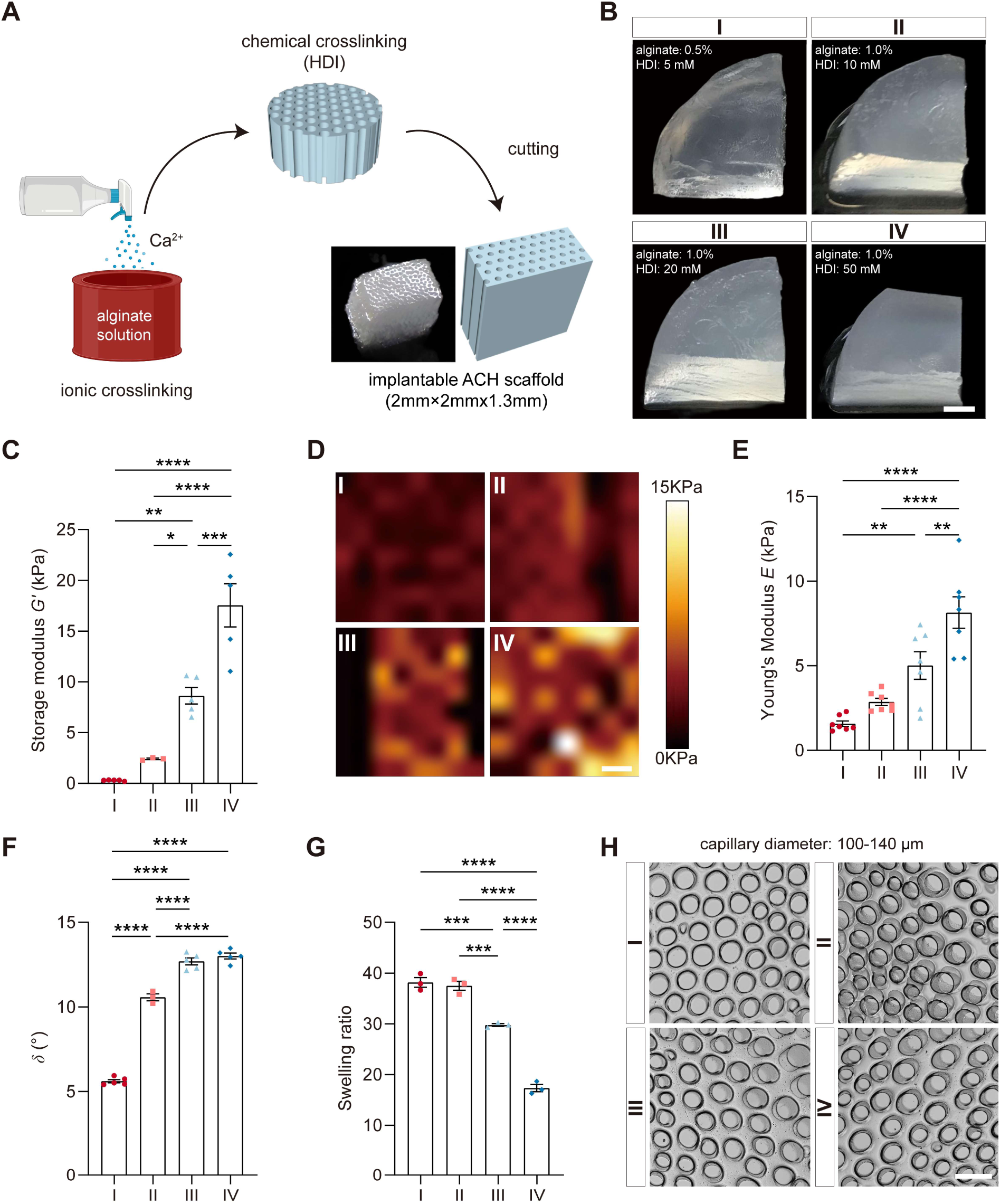
Mechanical and microstructural characteristics of stiffness varied ACHs. (A) A schematic diagram illustrating the fabrication process of ACHs. ACHs were produced by spraying Ca^2+^ cations into an alginate solution, followed by HDI chemical crosslinking for stabilization. (B) Representative pictures of each ACH block type with varying alginate and HDI concentrations. The ACH blocks showed increased transparency from type IV to I. Scale bar: 0.5 cm. (C) Storage modulus of all types of ACH blocks measured by rheological analysis in PBS at 37℃ (N=3-5). (D, E) Force maps and quantified apparent Young’s modulus on the surface parallel to capillaries in varied types of ACH cuboids, acquired from AFM measurements in DPBS at 37 °C (N=7). Invalid pixels in each force map are displayed by the color of the minimal value (black). Scale bar: 20 µm. (F) Stress-strain *δ* values for viscoelasticity characteristic of all types of ACH blocks (N=3-5). (G) Swelling ratio of all types of ACH blocks in PBS (N=3). (H) Brightfield images revealed the capillary microstructures of all types of ACH cuboids, approximately 120 µm. Scale bar: 250 µm. Statistical analysis was performed using a one-way ANOVA followed by a Tukey’s post hoc test (* *P* < 0.05, ** *P* < 0.01, *** *P* < 0.001, **** *P* < 0.001; error bars represent the standard error of the mean).

Consistent with prior AFM observations[19a], the Young’s modulus of the adult rat spinal cord falls within the range of a few hundred Pa, making type I the softest ACH approaching the host spinal cord stiffness while retaining its structure. All ACHs exhibited a viscoelastic nature dominated by elastic behaviors, as the phase-shift *δ* values from rheometric testing below 15° (Figure 1F). Given the sample’s storage modulus (*G’*) increases from type I to IV, the recorded loss modulus (*G’’*) increased even more distinctly, alluding to a slightly more balanced viscoelastic nature of those stiffer samples. Regarding stability, all ACH types released minimal alginate during a 4-week phosphate-buffered saline (PBS) immersion, indicating their non-biodegradable nature *in vitro*. However, the softer type I and II ACHs displayed greater weight-based swelling during the 4-week immersion period compared to type III or IV ACHs (Figure 1G). Structurally, the implantable ACH cuboids exhibited consistent capillary diameters of ∼120 µm and density of ∼40 capillaries/mm^2^ (Figure 1H, Figure S1, Supporting Information).

Having successfully fabricated a stiffness range of uniform microstructural ACHs closer to that of the rat spinal cord than we have previously used, we examined their response when implanted into rat cervical level 5 (C5) injured spinal cords (**Figure 2A**). Prior to implantation, all ACH scaffold types (dimensions 2×2×1.3 mm) underwent a coating process with poly-L-ornithine (PLO) and laminin. This coating was previously shown to neutralize the negative charge of the alginate in physiological conditions and enhance cell adhesion.^[5b]^ The PLO/laminin coating had no discernible impact on the apparent Young’s modulus measured by AFM (Figure S1, Supporting Information). The implantable ACH scaffolds were promptly placed into C5 lateral hemisections, effectively spanning the gap with the capillaries aligned parallel to the rostral-caudal axis (Figure S2, Supporting Information). Figure 2B illustrates the successful integration of all ACH scaffold types into the host spinal cord, with no overt degradation observed from the 4-week implantation period. Noteworthily, the less elastic type I ACH scaffolds (Figure 1F) were more prone to compressive forces of the host tissue and spine movement due to the activity of the animals, consequently leading to pressure on the scaffold and resulting in slight changes in scaffold size 4 weeks post-implantation (Figure 2B).

**Figure 2:**
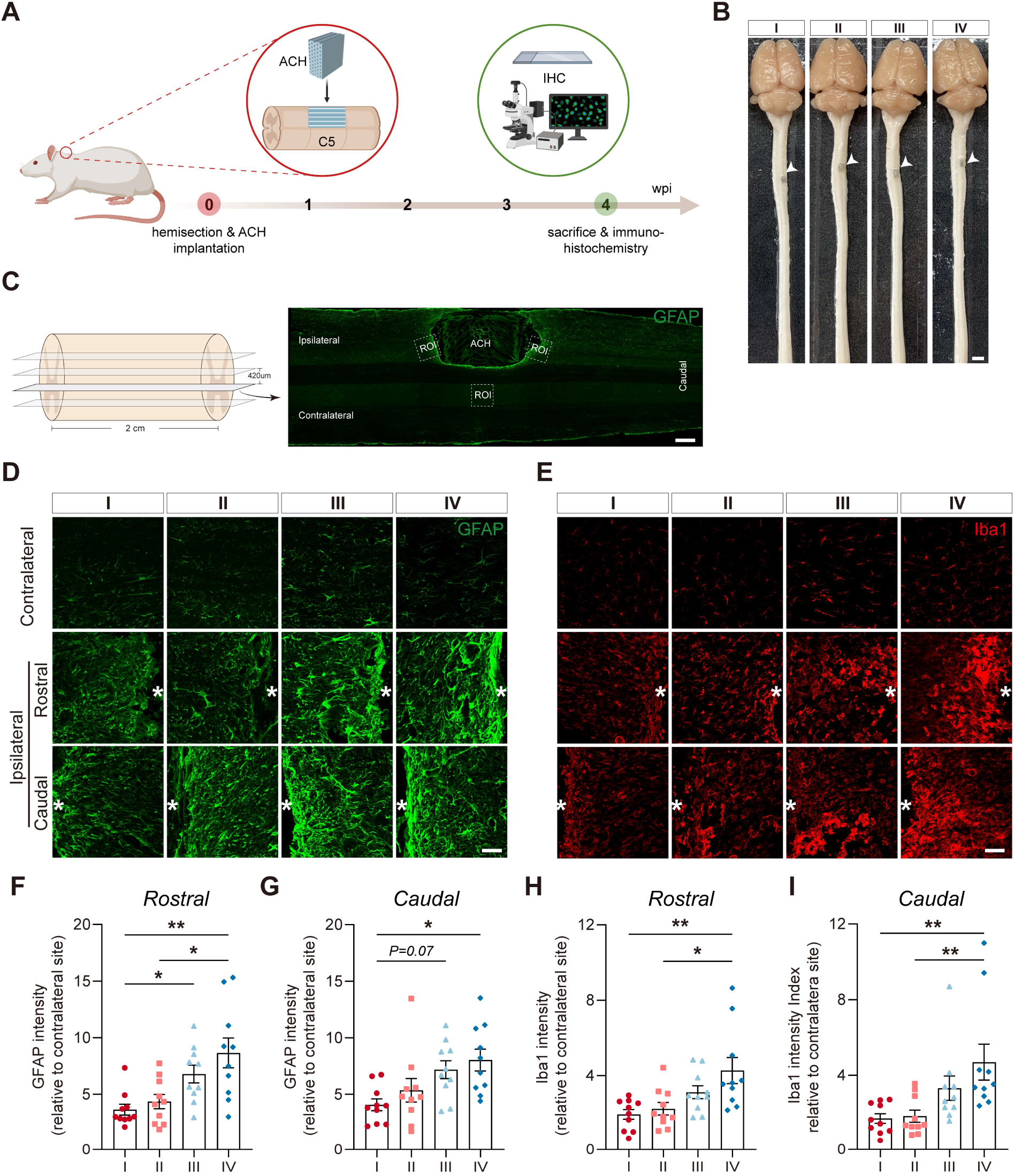
Greater foreign body reaction induced by stiffer ACH scaffolds. (A) A schematic diagram depicting the *in vivo* experimental setup to investigate the foreign body reaction to biomaterial stiffness. Following a cervical level 5 (C5) lateral hemisection (2 mm in length) of the rat spinal cord, varying stiffness ACH scaffolds were implanted into the lesion cavity with capillaries aligned along the rostral-caudal axis. After a 4-week survival period, animals were sacrificed for immunohistochemical analysis. (B) Representative images obtained during tissue processing demonstrates the successful integration of all ACH scaffolds into the host spinal cord without overt signs of cavitation or degradation. Scale bar: 2 cm. (C) Schematic diagram illustrating the quantitative analysis of immunolabeling for inflammation, with GFAP as an example. Sections were selected at 420 µm intervals. On each section, two regions of interest (ROIs) measuring 350 × 350 µm were chosen around ACH scaffolds, both rostrally and caudally on the ipsilateral side, in comparison to one ROI on the contralateral side. Scale bar: 500 µm. (D, E) Representative images showing the immunolabeling of astrocytes (GFAP) and microglia/macrophages (Iba1) within each ROI for each experimental group (white asterisks as the site of scaffolds), along with the quantification of intensity (F-I) on the ipsilateral side relative to the contralateral side (N=10). Scale bar: 100 µm. Statistical analysis was conducted using a Kruskal-Wallis test followed by a Dunn’s test (\**P* < 0.05, \*\**P* < 0.01, error bars: standard error of the mean).

Following SCI, the formation of a fibroglial scar restrains the lesion site from spreading secondary damage and repairs the spinal blood barrier, however, in the process of doing so it produces a physical and chemical barrier blocking neuroregeneration.[3] The foreign body reaction to biomaterials bridging the lesion site, adding to the fibroglial scar, poses a dual challenge by hindering both host-implant integration and neuroregeneration in the CNS.[6] We investigated the foreign body reaction triggered by stiffness-varied ACH scaffolds through immunolabeling astrocytes with glial fibrillary acidic protein (GFAP), microglia/macrophages with ionized calcium-binding adaptor molecule 1 (Iba1), and fibronectin (Figure 2C-E, Figure S3, Supporting Information). This observation strongly indicates a noteworthy astrogliosis and inflammatory reaction to the ACH scaffolds (ipsilateral), with the mildest reaction found in contact with type I scaffolds, and nearly doubling in contact with type IV scaffolds, indicating scaffold stiffness approaching the host spinal cord leads to less reaction (Figure 2D-I). This aligns with previous findings in the brain that have documented reduced neuroinflammatory responses when softer implants are utilized, as opposed to unphysiologically stiff ones.[8, 25] Nevertheless, the fibronectin area surrounding the scaffold exhibited minimal variation among the stiffness-varied groups at either end (Figure S3, Supporting Information), indicating the independent role of scaffold stiffness in fibronectin deposition elicited by the implanted scaffolds, which is in accordance with recent observations in the context of subcutaneous or peripheral nerve implants *in vivo*.[26] Hence, optimizing the host foreign body reaction might be achievable by closely mimicking the mechanical properties of biomaterials to match the native tissue of the CNS.

Glia make up the majority of the cellular population of the spinal cord and in response to injury reactive astrocytes can be polarized into pro-inflammatory (A1) or anti-inflammatory (A2) phenotypes.[27] Given the strong astrocytic differences observed between the ACH scaffold types and that mechanical stiffness can influence the polarization of astrocytes *in vitro*,[13] we investigated whether the scaffold stiffness affects the phenotype of A1 reactive astrocytes surrounding the scaffolds. We observed an increased complement 3 (C3)^+^GFAP^+^ region when in contact with type IV scaffolds compared to softer ones, indicating an enhanced presence of A1 astrocytes surrounding the stiffest ACH scaffold (**Figure 3A-B**).

**Figure 3:**
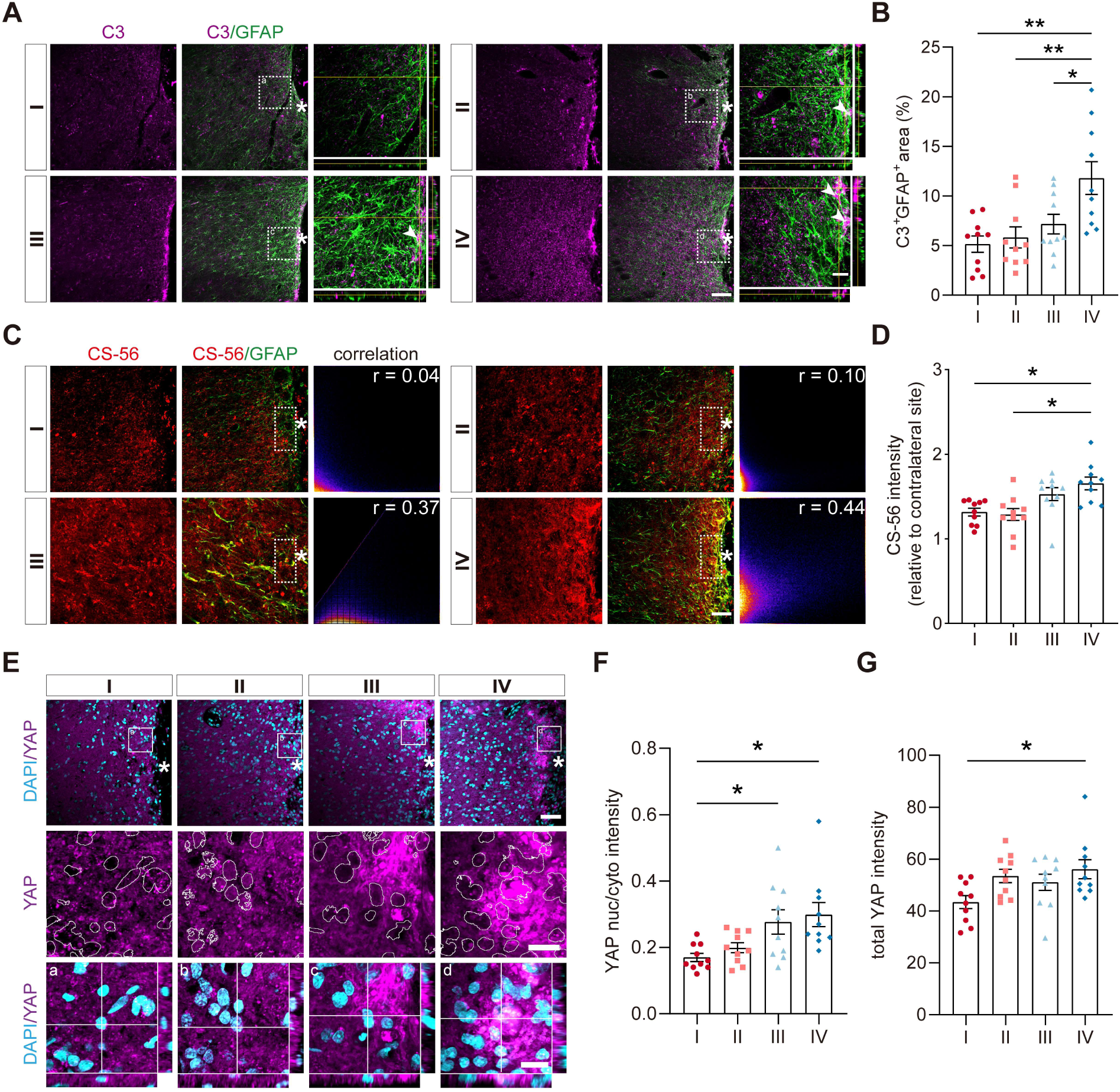
Increased inhibitory molecules are released and mechanosensing occurs in response to stiffer ACH scaffolds. (A) Co-immunolabeling of astrocytes (GFAP) and A1 astrocytes (C3 component) surrounding stiffness varied ACH scaffolds (white asterisks). Z-stack images on the right reveal spatial colocalization (indicated by white arrows). Scale bar: 100 µm (left); 30 µm (right). (B) Quantifying the colocalization area (C3^+^GFAP^+^) around scaffolds (N=10). (C) Co-immunolabeling of CSPG (CS-56) and GFAP around scaffolds (white asterisks) in each group, with accompanying intensity scatterplots on the right. Pearson coefficient r is displayed in the upper right corner. Scale bar: 20 µm. (D) Quantifying CS-56 intensity in regions surrounding ACH scaffolds on the ipsilateral side compared to the contralateral side (N=10). (E) Co-immunolabeling of YAP and nuclear DAPI in host tissue around varying stiffness ACH scaffolds (white asterisks). Higher magnification images in the two bottom rows emphasize YAP expression inside and outside the nucleus. Scale bar: 60 µm (top), 15 µm (bottom). (F, G) Quantifying nuc/cyto YAP and total YAP intensity around scaffolds in each group (N=10). Statistical analysis was conducted using a one-way ANOVA followed by a Tukey’s post hoc test for *B*, *F*, *G* and a Kruskal-Wallis test followed by a Dunn’s test for *C* (* *P* < 0.05, ** *P* < 0.01, error bars: standard error of the mean).

The ECM of the scar reinforced by the foreign body reaction forms a barrier that hinders neuroregeneration post-implantation. Chondroitin sulfate proteoglycans (CSPGs), a vital component of ECM, can also play a role in inhibiting the regenerative process.[3] Reactive astrocytes serve as one source of CSPG secretion following injury. A component of CSPGs, CS-56, was found elevated around the scaffold in comparison to the contralateral side. Similar to the inflammatory response, we observed a positive correlation between CS-56 expression and ACH scaffold stiffness, stiffer scaffolds resulted in greater CSPG deposition (Figure 3C-D). Additionally, the correlation coefficient r between CS-56 and GFAP was higher in the stiffer groups (Figure 3C), suggesting a greater degree of colocalization between CSPGs and astrocytes. Collectively, these findings imply that scaffold stiffness not only influences activation of astrocytes, marcrophages and microglia, but also their secretory capacity, aligning with previous *in vitro* observations.[13]

To investigate how the foreign body reaction was regulated by scaffold stiffness, we explored mechanotransduction, the translation of mechanical inputs into biochemical signals.[16] Among critical downstream mediators, the transcription factor YAP plays a pivotal role in mechanotransduction signaling originating from RhoA pathways or cytoskeletal tension.[7] When a cell detects mechanical cues related to stiffness, YAP undergoes translocation from the cytoplasm to the nucleus following a series of signaling events. This translocation subsequently leads to alterations in gene expression and cell behavior.^[17a, 28]^ In our current study, we observed an enhanced accumulation of YAP within the nuclei surrounding the stiffest ACH scaffolds when compared to the softest ones (Figure 3E-F). Moreover, the total intensity of YAP was highest in the stiffest group (Figure 3G), which had also shown heightened astrocyte proliferation (Figure 2D, F, G) and activation (Figure 3A-B) upon contact, previously shown to be activated by the RhoA pathway in SCI.[29] Together, these observations imply that the regulation of the foreign body reaction is contingent upon the ability of the cells surrounding the scaffold to sense stiffness, aligning with prior research conducted on various cell types *in vitro* and implant studies conducted *in vivo*.^[14a, 17c]^

A foreign body reaction to implants in the CNS often acts as a physical barrier blocking neuroregeneration. Hence, we hypothesized that when in contact with stiffer ACH scaffolds an increase in inflammatory cells and ECM deposition occurs leading to matrix buildup and subsequently increases this inhibitory barrier. To substantiate this, we examined host cell infiltration within the scaffolds. General nuclear 4’, 6-diamidino-2-phenylindole (DAPI) immunostaining revealed that host cells can infiltrate ACH scaffold capillaries from either end, with a preference for softer capillaries, as indicated by increased cell numbers and DAPI^+^ capillaries (**Figure 4 A-C**). Among these infiltrating cells, approximately 40% were microglia/macrophages (Iba1^+^) (Figure S4, Supporting Information). For the first time in implanted ACH scaffolds without additional interventions, a few host astrocytes were found to enter the capillaries of type I scaffolds aligned with axons, in contrast to the other three types (Figure 4D). Therefore, this finding indicates the weakest inhibitory barrier surrounds the softest ACH scaffolds augmenting cell infiltration.

**Figure 4:**
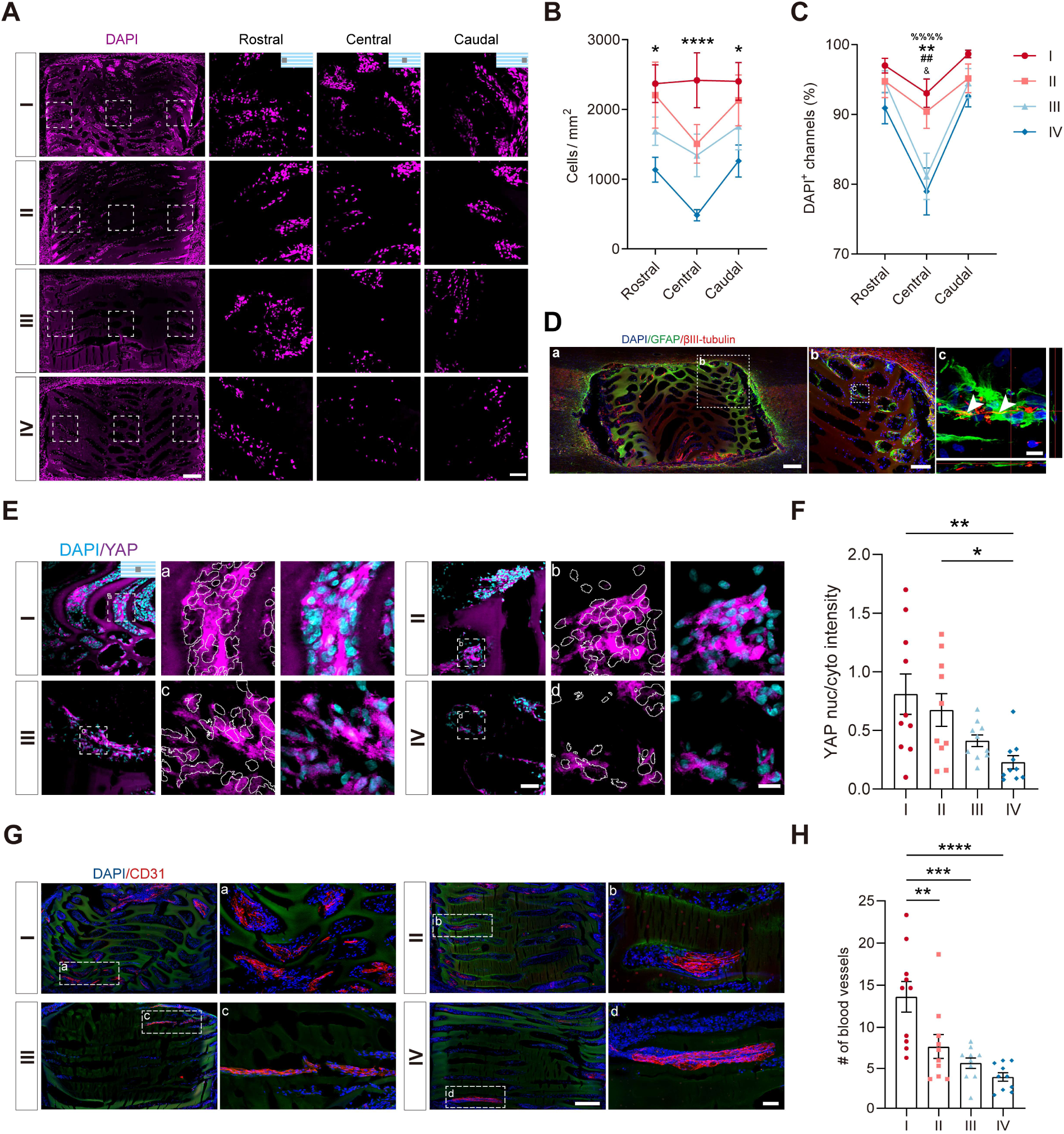
Increased cell filling of softest ACH scaffolds coincide with enhanced angiogenesis. (A) Immunolabeling of DAPI^+^ nuclei reveals cellular infiltration into capillaries in each group of implanted scaffolds. Higher magnification images in the three columns to the right correspond to the rostral, central, and caudal aspects of the scaffolds, as illustrated by the schematics in the first row. Scale bar: 200 µm (left); 50 µm (right). (B, C) Quantifying the number of infiltrated cells per square millimeter and the percentage of DAPI^+^ capillaries in each aspect of the scaffolds for each group (N=10). (D) Representative co-immunolabeling images of DAPI, GFAP, and axonal βIII-tubulin in type I ACH scaffold capillaries. Both an overview (a) and higher magnification images (b, c) show GFAP expression within the capillaries. Z-stack images (c) reveal colocalization between GFAP and βIII-tubulin (white arrows). Scale bar: 200 µm (a); 100 µm (b); 20 µm (c). (E) Co-immunolabeling of YAP and DAPI in the central aspect of scaffold capillaries for each group. Higher magnification images on the right highlight YAP expression within and outside the nuclei for each group. Scale bar: 50 µm (left); 15 µm (right). (F) Quantifying nuc/cyto YAP intensity within the central capillaries of varied scaffolds (N=10). (G) Immunolabeling of regrowing blood vessels (CD31) within scaffold capillaries, with higher magnification images on the right for each group. Scale bar: 200 µm (left); 50 µm (right). (H) Quantifying the number of blood vessels within scaffold capillaries (N=10). Statistical analysis was performed by using a two-way ANOVA followed by a Tukey’s post hoc for *B*, *C* (* *P_I vs IV_* < 0.05, ** *P_I vs IV_* < 0.01, **** *P_I vs IV_* < 0.0001, ^&^ *P_II vs IV_* < 0.05, ^##^ *P_I vs III_* < 0.01, ^%%%%^ *P_I vs II_* < 0.0001, error bars: standard error of the mean) and a one-way ANOVA followed by a Tukey’s post hoc test for *F*, *H* (* *P* < 0.05, ** *P* < 0.01, *** *P* < 0.001, **** *P* < 0.0001, error bars: standard error of the mean).

In type II-IV ACH scaffolds, there were noticeably fewer cells within the central region compared to either entry region. Conversely, infiltrating cells remained relatively consistent throughout the entire type I scaffold (Figure 4B), suggesting that the softer scaffold also promotes cell migration, aligning with previous *in vitro* findings.[30] Nuclear YAP expression has been shown to improve cell motility by allowing adaptive cytoskeletal remodeling and limiting focal adhesions,[31] which matches the finding of type I ACH scaffolds exhibiting the greatest nuclear YAP in the central region compared to the other types (Figure 4E-F). Moreover, no significant differences were observed in YAP nuclear translocation among groups at both the rostral and caudal entries (Figure S5, Supporting Information). Although there was a trend towards lower YAP in the stiffer scaffolds which had less cell infiltration. Consequently, we observed increased migration and/or proliferation of cells within the soft scaffolds leading to enhanced YAP signaling, conversely when the foreign body reaction is initiated by stiff scaffolds enhanced cell migration and/or proliferation leads to greater YAP activation.

Angiogenesis plays a crucial role in providing trophic support for infiltrating cells as well as neural regeneration after scaffold implantation.[32] It primarily relies on the proliferation of endothelial cells, a process potentially influenced by stiffness-dependent cell infiltration. Furthermore, recent studies have indicated that both pathological and physiological angiogenesis processes are influenced by signals related to environmental stiffness.[17b, 33] Subsequently, we conducted an analysis of angiogenesis within the capillaries by immunolabeling CD31, also known as platelet endothelial cell adhesion molecule (PECAM-1). After implantation, the coating of laminin contributes to the formation of a basement membrane for recruiting endothelial cells.[34] Remarkably, among groups with varying stiffness, type I ACH scaffolds exhibited significantly larger CD31-positive areas compared to type III or IV scaffolds (Figure 4G). Additionally, manual counting of CD31^+^ blood vessels throughout the entire scaffold showed a significant decrease with increased scaffold stiffness (Figure 4H). This indicates that scaffold stiffness can have a profound impact on angiogenesis within the implanted scaffold. A prior study has reported that YAP signaling drives proliferation and re-arrangement of endothelial cells in developing vessels.[17b] Similarly, the increased angiogenesis within the center of the scaffolds we observed here was associated with heightened cell migration and enhanced YAP nuclear translocation, likely triggered by the softer environment.

Considering the improved ACH scaffold microenvironment resulting from the limitation of the foreign body reaction and enhanced internal migration and/or proliferation of infiltrating cells, axonal regeneration within the implanted scaffold may also be influenced. Additionally, previous *in vitro* studies have highlighted the role of stiffness in regulating axonal growth patterns, neurite extension, and branching.[9, 15a] In particular, a recent *in vitro* study has shown that a softer natural polymer scaffold, similar to the type I scaffold in our study, collectively modulates primary astrocyte behavior and enhances axonal growth of seeded dorsal root ganglion neurons.[35] We then investigated the impact of scaffold stiffness on axonal regeneration by visualizing the regrowth of both descending and ascending axons within ACH scaffold capillaries with a general mature neuronal marker βIII tubulin. Not surprisingly, both ascending and descending axons entered the implanted scaffold regardless of its stiffness (**Figure 5A**). Generally, the regrowing axons formed clustered, branched bundles within the capillaries. Quantitatively, the regrowth of axons was notably higher in type I ACH capillaries compared to type IV at both entries of the scaffold (100 µm away from the scaffold’s edge) (Figure 5B). Moreover, it was observed that the softer scaffolds housed a greater number of axons within their entire capillaries, indicating that the softer environment attracted a higher number of regrowing axons (Figure 5C). Additionally, 5-hydroxytryptamine (5-HT) immunolabeled descending raphespinal serotonergic axons were most abundant within the rostral entry of type I ACH scaffolds, although they decreased towards the caudal side in all groups (Figure 5D-F). Together, these findings corroborate a prior study that demonstrated a growth preference for softer environments by retinal ganglia axons.^[15b]^ Here, we present direct evidence of the relationship between biomaterial stiffness and axonal regeneration in *in vivo* CNS tissue engineering.

**Figure 5:**
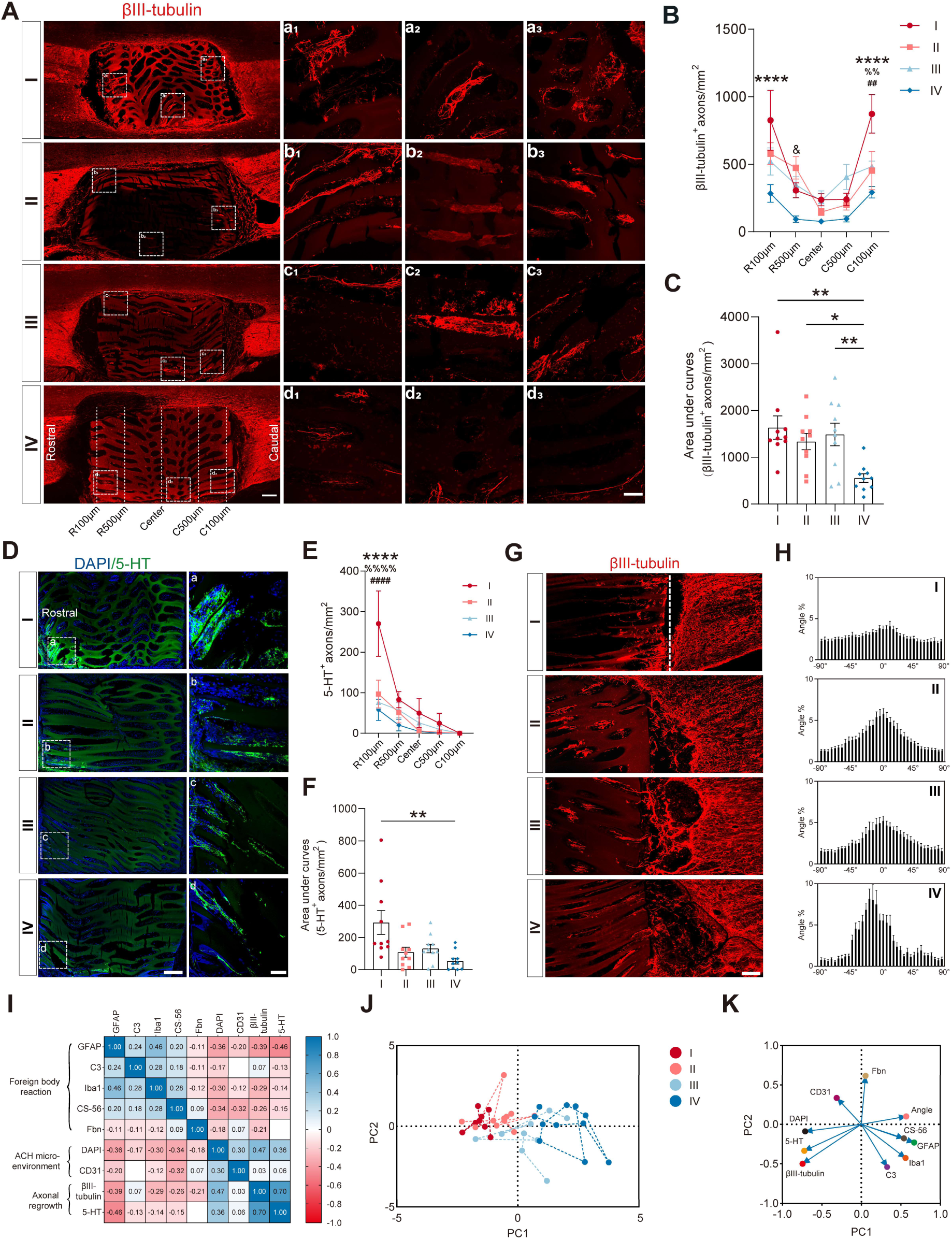
Softer ACH scaffolds enhance axonal growth, but not directionality, influenced by multiple foreign body reaction factors. (A) Immunolabeling of both descending and ascending axonal regrowth (βIII-tubulin) within varying stiffness ACH scaffolds. Higher magnification images in the three right columns were selected from the white dotted boxes on the left for each group. Scale bar: 200 µm (left); 100 µm (right). (B, C) Quantifying the density of βIII-tubulin positive axons per square millimeter at various distances along the virtual lines (white dotted lines in *A*) and the area under curves for each group (N=10). (D) Immunolabeling of serotonergic axonal regrowth (5-HT) within ACH scaffolds of varying stiffness. Higher magnification images in the right column are provided for each group. Scale bar: 200 µm (left); 100 µm (right). (E, F) Quantifying the density of 5-HT positive axons per square millimeter at various distances along the virtual lines and the area under curves for each group (N=10). (G) βIII-tubulin labeled images highlight the orientation of regrowing axons within scaffold entries in each group, with a white dotted line indicating the host-implant interface. Scale bar: 100 µm. (H) Quantification of the angle distribution of regrowing axons within scaffold capillaries in each group. ±90° represents vertically oriented axons, while 0° represents horizontally oriented axons. (I) Correlation map combining all markers for distinct biological activities. (J) Principal component analysis of the quantified markers (GFAP, Iba1, C3, CS-56, Fbn, DAPI, CD31, βIII-tubulin, 5-HT, Angle). Softer scaffolds (type I and II) are grouped on the left side, gradually transitioning to the stiffer scaffolds (type III and IV) on the right. (K) Eigen loadings indicate the relationship between markers and PC1. Statistical analysis was conducted by using a two-way ANOVA followed by a Tukey’s post hoc test for *B*, *E* (**** *P*_I vs IV_ < 0.0001, ^&^ *P*_II vs IV_ < 0.05, ^##^ *P*_I vs III_ < 0.01, ^####^ *P*_I vs III_ < 0.0001, ^%%^ *P*_I vs II_ < 0.01, ^%%%%^ *P*_I vs II_ < 0.0001, error bars: standard error of the mean) and a Kruskal-Wallis test followed by a Dunn’s test for *C*, *F* (* *P* < 0.05, ** *P* < 0.01, error bars: standard error of the mean).

Remarkably, when compared to their entries, we observed a significant reduction in the number of regrowing axons towards the center of the scaffolds. No significant differences were observed among the various groups (Figure 5B). This suggests that while softer ACH scaffolds attract more axons, they do not always promote further extension. This observation aligns with a previous study that reported slow and incoherent axonal growth in a soft environment *in vitro*.[15b] To examine this, we conducted measurements of the directional axonal growth within the scaffold capillaries. We utilized AngleJ,[36] a plugin of ImageJ, to categorize the various angle ranges between ±90° of regrowing axons, with 0° representing the horizontal direction parallel to the rostral-caudal orientation of the capillaries (Figure S6, Supporting Information). Interestingly, we observed a uniform distribution of axonal angles in type I ACH scaffolds, suggesting that regrowing axons exhibited equal growth in every direction (Figure 5G-H). In contrast, for the other types of scaffolds, a clear bell-shaped distribution of axon angles was evident, indicating a higher prevalence of a rostral-caudal growth direction (Figure 5G-H). Furthermore, axons exhibiting rostral-caudal orientations (±10° or ±20°) were more abundant within the stiffest ACH capillaries compared to the softer ones (Figure S6, Supporting Information). Hence, the lack of axon extension within the softer scaffold may be attributed to the non-linear growth of axons.

Nevertheless, anisotropic capillaries are purposefully designed to physically guide axonal growth in a specific direction, limiting tortuous axonal growth typically observed for isotropic injectable biomaterials.[37] Here, we showed that softening the ACH leads to nonlinear axonal regrowth only in the softest type I scaffolds, while the other types all showed a prevalence of linear growth along the rostral-caudal axis (Figure 5G-H). It is known that neurons sense mechanical stiffness and grow towards softer substrates, once there slow their growth and splay their axons in response.[15b] Further, it has been discovered that axonal splaying can be influenced by vascular density,[38] which is significantly greater in type I ACH (Figure 4G-H). This underscores the requirement for a balance between making the biomaterial scaffold soft, akin to the original tissue, to avoid the foreign body reaction, but also providing directional guidance cues, when lacking soluble guidance molecules. Furthermore, it is worth noting that the softest ACH capillaries bend more frequently after implantation possibly due to the combined swelling of the ACH and compressive pressures from the surrounding host tissue leading to instability. In our study, 5 of 10 implanted ACH scaffolds from the type I group exhibited capillaries which deviate from the rostral-caudal axis by 45° (Figure 5A, D), whereas the other scaffold types did not deviate to such an extent.

We observed that most of the foreign body reaction markers exhibited negative correlations with markers for scaffold microenvironment and axonal regrowth. Conversely, an improved scaffold microenvironment was associated with greater axonal regrowth (Figure 5I). These findings were further validated through a principal component analysis, which indicated that softer scaffolds (type I and II) were aligned with a favorable scaffold microenvironment (DAPI and CD31) and axonal regrowth markers (βIII Tubulin and 5-HT) while being less associated with most of the foreign body reaction markers (GFAP, Iba1, C3, CS-56, and Fbn) (Figure 5J-K). Collectively, our results suggest that enhanced axonal regrowth in softer scaffolds is influenced not only by the collective reduction of the host foreign body reaction but also by improvements in the scaffold microenvironment. Allowing more cells to infiltrate into softer scaffolds benefits angiogenesis and axonal ingrowth. However, our own personal observation is that doubling the amount of capillaries does not affect stiffness or cell infiltration.[39] This work was done with non-degrading scaffolds, it will be of interest to examine over time the change in stiffness with degrading scaffolds and the role this plays in the foreign body response and regeneration.

There is a growing number of studies aimed to explore the mechanical properties of CNS tissue as an important indicator in different conditions, including during development or tissue repair.[18, 23, 40] However, there has been a lack of investigation into the mechanical properties of host tissue in response to *in vivo* implanted scaffolds. The stiffness of CNS tissue post-injury has been linked to glial cell activation and the deposition of ECM components.[19b] Therefore, based upon this observation, it is conceivable that scaffold stiffness would influence the mechanical properties of the host tissue in response to implanted scaffolds. To investigate this hypothesis, we conducted micro-indentation AFM measurements on host-implant slices taken 4 weeks after implantation (**Figure 6A-B**). All slices were collected in artificial cerebrospinal fluid (aCSF) to maintain physiological properties (Figure 6A). In line with prior AFM observations^[19a]^, the apparent Young’s modulus values (*E*) for the uninjured spinal cord (R1/R1’ regions located approximately 4 mm from the scaffold) ranged from 10 to 300 Pa, with a median value of 75 Pa.

**Figure 6:**
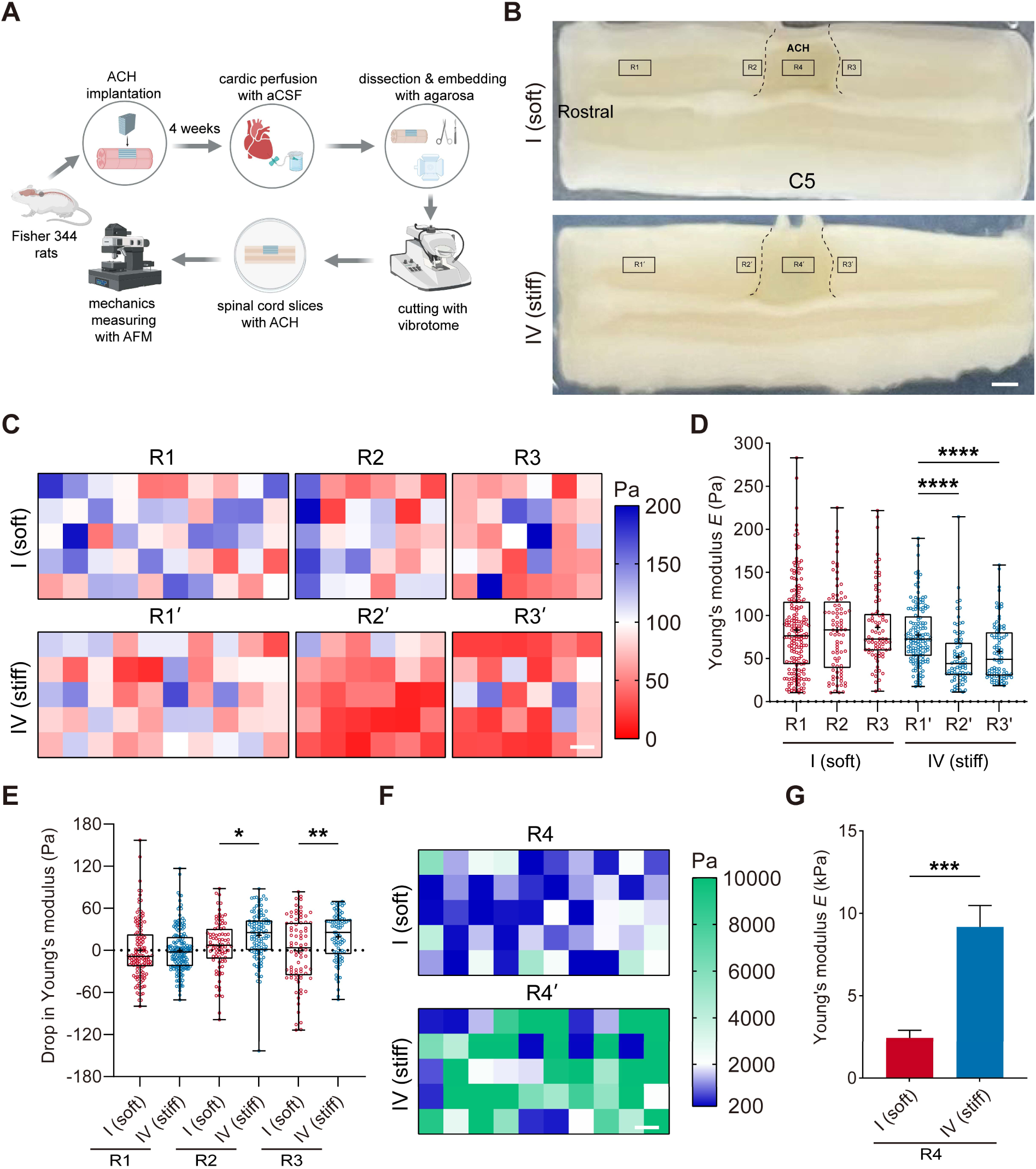
The stiffest scaffold softens surrounding host tissue four weeks post-implantation. (A) A schematic diagram of the *in vivo/ex vivo* experimental setup for investigating the mechanical impact of biomaterial stiffness. Four weeks after ACH scaffold implantation, animals were transcardially perfused with slicing artificial cerebral spinal fluid (aCSF). The spinal cord containing the ACH scaffold was then dissected and sliced for AFM measurements in slicing aCSF at 37°C. (B) Representative images of host-implant slices for each group. For AFM measurements, 4 distinct regions of interest (ROIs) were selected for each animal. R1/1’ (500×1000 µm) is rostrally located on the spinal cord, 3 mm away from the scaffold. R2/2’ and R3/3’ (500×500 µm) are placed on the spinal cord adjacent to the scaffold at rostral and caudal sides, respectively. R4/4’ (500×1000 µm) is positioned in the center of the scaffold. Scale bar: 700 µm. (C) Stiffness maps and (D) quantification of apparent Young’s modulus for the R1/1’, R2/2’, and R3/3’ regions (N=3). Scale bar: 100 µm. (E) A drop in Young’s modulus represents the apparent Young’s modulus values in each region relative to the values of the R1/1’ region for each animal. (F) Stiffness maps and (G) quantification of apparent Young’s modulus for the R4/4’ regions (N=3). Scale bar: 100 µm. Statistical analysis was performed using a Mann-Whitney test (* *P* < 0.05, ** *P* < 0.01, *** *P* < 0.001, **** *P* < 0.0001, error bars: standard error of the mean).

**Figure 7:**
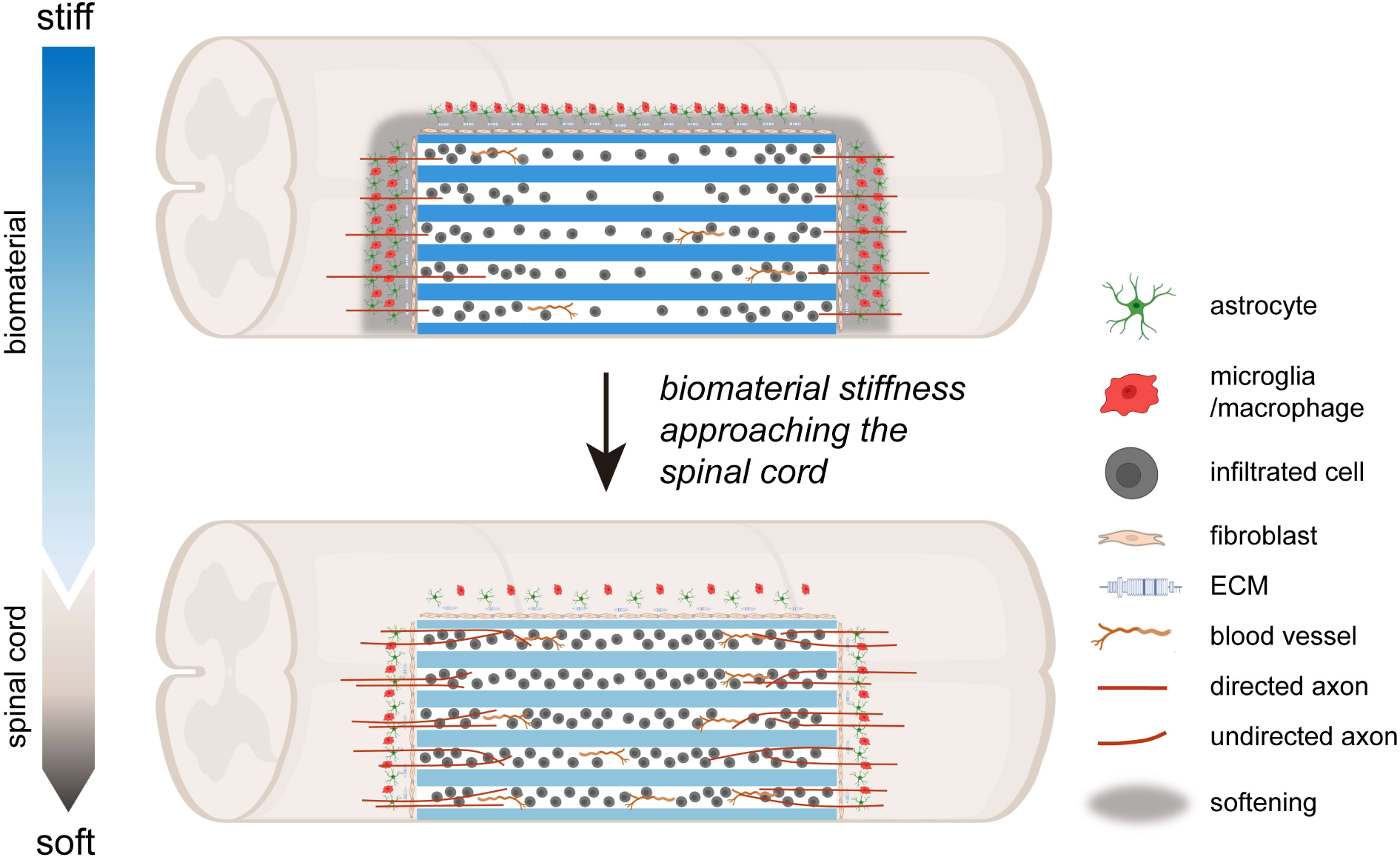
Biomaterial scaffold stiffness influences the foreign body reaction, tissue stiffness, angiogenesis and neuroregeneration in spinal cord injury. The proximity of biomaterial stiffness to native tissue exhibits a remarkable reduction in foreign body reactions, encompassing astrocyte and microglia/macrophage activation, after implantation. This enhanced foreign body reaction surrounding the biomaterial coincides with host tissue softening. In contrast, the enhanced cellular infiltration and angiogenesis triggered by attenuated foreign body reaction foster axonal ingrowth, although needs to be optimized to avoid undirected growth.

Implanting type I (*E* 1kPa) or type IV (*E* 9 kPa) ACH scaffolds for four weeks did not alter the stiffness of the remote uninjured spinal cord tissue. However, in the regions of the spinal cord adjacent to the scaffolds (R2/R2’ or R3/R3’ regions), we notably observed a significant reduction in apparent Young’s modulus (∼30 Pa) compared to the uninjured region in the type IV group, whereas the type I group did not exhibit this effect (Figure 6C-E, Figure S7, Supporting Information). Additionally, the difference in apparent Young’s modulus within the two types of implanted scaffolds remained consistent with what was observed prior to implantation (Figure 6F-G), suggesting that implantation did not alter the mechanical properties of the ACH scaffold. It also implies that the degradation of both types of scaffolds did not appear 4 weeks after implantation. Together, these findings indicate that contact with the stiffest ACH scaffold softens the adjacent host tissue, whereas the softest ACH scaffold does not have this effect. The distinct foreign body reaction caused by varying scaffold stiffness may underlie this mechanical effect, as indicated by the previous study regarding spinal stab injury.[19b]

## 3. Conclusion

Here, we have shown approaching the stiffness of the biomaterial scaffold to that of the original tissue can minimize the host foreign body reaction and retain the mechanical properties of the host tissue, while improving the scaffold microenvironment. This ultimately leads to enhanced axonal regeneration and angiogenesis through the scaffold. However, too closely matching spinal stiffness can result in scaffold instability and non-linear axonal regrowth within the scaffold, similar to issues faced by softer isotropic injectable biomaterials.[37] Therefore, a more balanced approach for a softer (1.5-3 kPa in elastic modulus) but not the softest scaffold allows for the retention of linear physical guidance without the enhanced foreign body reaction. These findings underscore the complex role of stiffness in biomaterial scaffold design for tissue engineering and contribute to a better understanding of the interplay between the scaffold mechanical properties and host cellular responses. This knowledge can already be applied to scaffolds in clinical trials to minimize patient foreign body reactions and improve spinal integration, in hopes of increasing SCI patient recovery.[41]

## 4. Experimental Methods

*Materials*: PRONOVA UP MVG (1003-03) medical-grade alginate was sourced from NovaMatrix (Sandvika, Norway). This sodium alginate product is characterized by medium viscosity (η > 200 mPas, Mw = 75-200 kDa), with a guluronic acid content of ≥ 60% (w/w) and a protein content of ≤ 0.3% (w/w). The bivalent inorganic salt Ca(NO_3_)_2_·4H_2_O (purity ≥ 99.0% (w/w)) and hexamethylene diisocyanate (HDI) (purity ≥ 99% (w/w)) were procured from Sigma-Aldrich (St. Louis, USA). Hydrochloric acid solution (1 M) was acquired from VWR international S.A.S. (Fontenay-sous-Bois, France). Ethanol (p.a.) (purity ≥ 99.8% (v/v)), and acetone (p.a.) (purity ≥ 99.5% (v/v)), were supplied by Sigma Aldrich (St. Louis, USA). Nitric acid (65% (v/v)) was sourced from Fisher Scientific UK (Loughborough, UK).

### Fabrication of alginate anisotropic capillary hydrogels (ACHs)

Medical-grade sodium alginate was dissolved in purified water overnight with stirring at concentrations of 0.5% or 1.0% (w/w). The alginate sols (65 g) were poured into nitric acid-oxidized aluminum caps and allowed to rest overnight. To produce anisotropic capillaries, the gels were covered with 1 M Ca(NO_3_)_2_ solution by utilizing a pressure nebulizer. After a minimal gelation period of 48 hours, the gels were washed with purified water to remove excess metal salt solution. For chemical cross-linking, the gel block underwent dehydration in ascending concentrations of acetone (25%, 50%, 75%, 100%), followed by dry acetone containing 5-50 mM HDI. The fixed gel blocks were then immersed in 0.1 M HCl to exchange ions followed with purified water until pH neutrality was obtained. To fabricate stiffness-varied ACH types, different concentrations of alginate and HDI were utilized: type I (alginate, 0.5%; HDI, 5 mM), type II (alginate, 1.0%; HDI, 10 mM), type III (alginate, 1.0%; HDI, 20 mM), type IV (alginate, 1.0%; HDI, 50 mM). For AFM measurements and *in vivo* experiments, gel blocks were cut into cuboid shapes (2×2×1.3 mm) with a vibratome (Leica VT1000S, Leica Biosystems GmbH). Scaffolds were then precisely inspected under a light microscope (Olympus BX53; Olympus Life Sciences) to ensure evenly distributed and oriented capillaries. For rheometry testing, whole gels were cut in slices of 3 mm. The resulting ACH samples were stored in 70% ethanol for sterilization.

### Rheological measurements

To assess the bulk mechanical properties of ACHs, 3-5 ethanol-preserved bandsaw-cut slices (3 mm in thickness) from each type underwent 3h immersion in PBS. Subsequent mechanical analysis employed a TA Instruments AR 2000 Rheometer. A frequency sweep (0.1-10 Hz, 100 µNm oscillatory torque) was executed utilizing a 40 mm stainless steel, cross-hatched plate with a solvent trap as the top geometry. Simultaneously, a Peltier element, covered by a stainless-steel plate featuring a 40 mm cross-hatched surface, served as the bottom geometry. Calibration of the measurement gap and geometry inertia preceded the measurements conducted at 37 °C. Following the complete cross-linking procedure, the sample disks exhibited a reduced thickness and diameter compared to their initial state after bandsaw cutting. Subsequently, as the gels were compressed and indented by the rough geometry surface, the measurement gap was meticulously adjusted in a range between 800 to 1100 µm. Given the diverse mechanical rigidity of the tested sample disks, the normal force was systematically varied between 0.2 and 2 N to mitigate potential issues, such as excessive syneresis and complete compression of softer ACHs. Storage Modulus *G’* and stress-strain *δ* values were obtained for an angular shearing frequency of 1 Hz from each frequency sweep. The recorded *δ* values indicate the viscoelastic nature of the samples, as a bigger phase-shift in the strain response of the sample indicates a more fluid-like behavior.[42] To preserve the structure of the hydrogel samples during the frequency sweep, the oscillatory torque and stress were orders of magnitude lower than those that would be present beyond the linear viscoelastic region in stress-sweep testing of identical samples.

### AFM measurements

To characterize the surface stiffness of varied ACH scaffolds, 7-8 ethanol-preserved scaffolds from each type underwent 3h immersion in Dulbecco’s phosphate-buffered saline (DPBS). Subsequent micro-indentation experiments were performed with a JPK Nanowizard (Life science, JPK Instruments, Bruker Nano GmbH) placed on a motorized fluorescence microscope (Olympus IX81, Olympus Life Sciences). As an indentation entity, rectangle silicon cantilevers attached with a 20 µm diameter colloidal particle tip at the very end part (CP-CONT-BSG-C, Bruker Nano GmbH) were used. Cantilever spring constants were calibrated by using the thermal noise method in the measuring solution at 37°C implemented by the AFM software (JPK SPM). A CCD camera (The Imaging Source) was mounted by side of the fluorescent microscopy to align and monitor the position of the cantilever over defined regions of the scaffold surface.

All measurements were performed at 37 °C in DPBS buffer using a set petri dish heater (JPK Instruments). Scaffold surfaces parallel to the capillary were approached by the cantilever tips. Force-distance curves were taken with an approach speed of 1 µm/s and a set force of 10-20 nN maximally. Three areas (100×100 µm) of the sample surface were investigated in a lateral-scanning mode by recording 64 measure points approximately every 12.5 µm. The apparent Young’s modulus *E* was calculated from the Hertz fit of the force curve based on a spherically shaped indenter with the JPK software 4.0.x. The subsequent equation 1 displays apparent Young’s Modulus *E* of a sample, which is indented by a spherical probe of radius *R_C_*. The present force *F* at several indention depths *δ* is monitored. The value of the Poisson ratio, denoted by *v*, is 0.5.

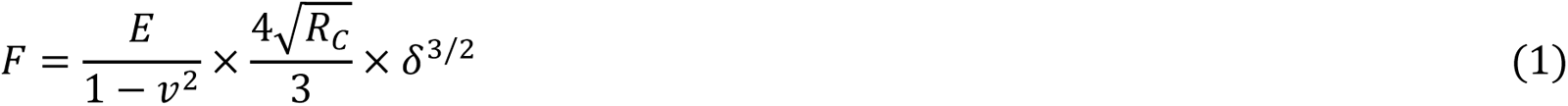

### Swelling rate

Three samples (1-2 g) from each ACH type were immersed in PBS overnight. Afterward, excess liquid was gently removed with a paper towel. The samples were then placed in weighed centrifugation tubes. Freeze-drying was performed with loosely tightened screw caps overnight at 4 to 8 mbar. The tubes were then re-weighed, and swelling rate was calculated using equation 2, with *w_s_* as the weight of the sample in its swollen state and *w_d_* as its weight after freeze-drying.

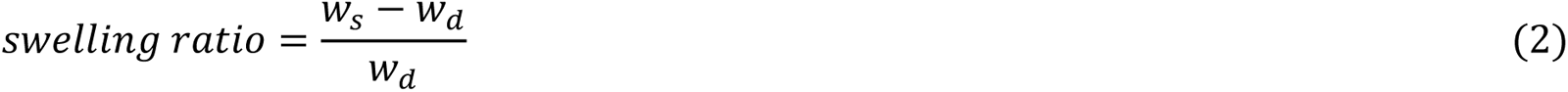

### Structural characterization

To analyze the capillary structure, 6 slices (400 µm in thickness) were obtained from each type of ACH using a vibratome and examined under brightfield. Ten images were captured from each type at a magnification of 10× using a fluorescent microscope. Capillary density and diameter were quantified using ImageJ to assess the structural characteristics of each type.

### Surface coating before in vivo implantation

All ACH scaffolds used *in vivo* were coated with PLO and laminin.[5b] One day before coating, scaffolds were transferred into DPBS for washing and stored overnight. Afterwards, PLO (Sigma Aldrich) was applied to the scaffolds at a concentration of 0.5 mg/mL in ice-cold dH_2_O and allowed to incubate overnight at 37°C with 5% CO_2_. The scaffolds were then washed twice in sterile DPBS for 30 minutes each on an orbital shaker (Neo Lab) at 75 rpm. Next, laminin was applied to the scaffolds at a concentration of 10 µg/mL in sterile DPBS and allowed to incubate for 2 hours at 37°C with 5% CO_2_. The PLO/laminin-coated ACH scaffolds were then rinsed twice for 30 minutes each in DPBS and stored in sterile DPBS at 4°C until further use.

### Animals

All animal experiments were conducted following national guidelines for animal care under the European Union Directive (2010/63/EU) and approved by the local governing institute (Regierungspräsidium Karlsruhe, G-195/19). Wild-type adult female Fischer-344 rats (> 180 g, 10-12 weeks old) were obtained from Janvier labs (Strain: F344/HanZtmRj) and utilized for all *in vivo* studies. Rats were housed in groups of 4-5 animals/cage on a 12/12-hour light/dark cycle with free access to food and water (*ad libitum*). Animal well-being, as well as temperature (20 ± 1°C) and humidity (45-65%), were checked daily by trained staff.

### Surgical procedures

As previously reported, all rats underwent a unilateral (right-side) spinal cord hemisection at C5 segment.[4b, 5] Following the hemisection, different types (I to IV) of ACH scaffolds were immediately implanted into the lesion cavity. Initially, the animals were administered with a mixture of ketamine (62.5 mg/kg), xylazine (3.175 mg/kg), and acepromazine (0.625 mg/kg) in 0.9% sterile saline via intraperitoneal (i.p.) injection to achieve deep anesthesia. The spinal column was exposed following a skin incision, and a laminectomy was performed. Thereafter, a rostro-caudal incision along the spinal midline was made into the dura with a scalpel to expose the spinal cord. To create a lesion, a unilateral block (2 mm in length) of the spinal cord tissue was carefully removed. ACH scaffolds (2×2×1.3 mm) were then directly implanted into the lesion cavity with their capillaries in a rostro-caudal direction. Following implantation, the dura was covered with a thin dried agarose film and sealed with tissue glue (fibrinogen (100 mg/ml) + thrombin (400 U/ml), Sigma Aldrich). The paravertebral muscle layers were readjusted, sutured, and the skin was stapled. To prevent surgery-related dehydration, each animal received a subcutaneous (s.c.) injection of 1 ml Ringer solution. To alleviate acute pain, all rats received s.c. injections of burphrenophin (0.03 mg/kg in sterile 0.9% saline, Reckitt Benckiser) and ampicillin (50 mg/kg in sterile 0.9% saline, Ratiopharm) to prevent wound infection during the first 2 days after surgery. To promote recovery, the animals were given a high-caloric drink (Fresubin, Fresenius Kabi) up to 3 times per day until their body weight stabilized. One animal implanted with type III ACH scaffold died during the surgical procedure and were then excluded.

### Tissue cryo-processing

Rats were euthanized four weeks after the surgery for immunohistochemical analysis. A lethal dose of a ketamine (62.5 mg/kg), xylazine (3.175 mg/kg), and acepromazine (0.625 mg/kg) mixture, diluted in sterile 0.9% saline, was administered intraperitoneally. The animals were then subjected to transcardial perfusion using 0.9% saline and followed with 4% paraformaldehyde (PFA)/0.1 M phosphate buffer (PB). The brains and spinal cords were carefully dissected and post-fixed for one hour in 4% PFA/0.1 M PB at room temperature. Thereafter, the tissues were cryoprotected at 4°C for at least two days in 30% sucrose/0.1 M PB. The cervical spinal cord was embedded in Tissue-Tek O.C.T.™ compound (Sakura) and horizontally cut into 30 µm thick sections using a cryostat (Hyrax C60, Zeiss). The tissue slices were mounted serially and directly onto glass slides, with every 4-5 slices on each slide. Slides were stored at -80°C until further use. ACH scaffolds detached from the spinal cords during dissection following perfusion in two animals (one implanted with type I and another with type IV ACH scaffold), who were excluded afterward.

### Immunohistochemistry

For the immunohistochemical analysis, sets of serial tissue slices (every 14^th^ slice, totaling 4-5 slices for one serial set) combining the cervical spinal cord and scaffold were utilized. The tissue slices were thawed and air-dried for an hour at room temperature and surrounded with a liquid blocker. The samples were then washed three times for 20 minutes each in tris-buffered saline (TBS) and incubated at room temperature with TBS + 0.25% Triton-X100 + 5% donkey serum for 2.5 hours to permeabilize the tissue and prevent unspecific antibody binding. Primary antibodies were diluted in TBS + 0.25% Triton-X100 + 5% donkey serum and incubated overnight at 4°C in humid conditions (Table S1, Supporting Information).

Subsequently, the tissue sections were washed three times for 20 minutes each at room temperature in TBS + 1% donkey serum. Secondary antibodies and DAPI were diluted in TBS + 1% donkey serum, and samples were incubated in the dark for 2.5 hours at room temperature (Table S1, Supporting Information). Lastly, the samples were washed three times for 20 minutes each in TBS, air-dried for 20 minutes at room temperature, and covered with Fluoromount-G.

### Quantification for immunostaining

For quantification, one set of serial tissue slices per animal were used. For each maker, images were taken with identical parameters (exposure time, gain, and offset) using a fluorescent microscope (Olympus BX53; Olympus Life Sciences) or a confocal fluorescent microscope (Olympus FV1000; Olympus Life Sciences). To quantify the host response and YAP signals, regions of interest (ROIs, 350×350 µm) were delineated from the area adjacent to the ACH scaffold (host-implant interface) on both the rostral and caudal sides on the ipsilateral site, as well as a single one on the corresponding contralateral site. Using ImageJ, the intensity or proportional above-threshold area of each ROI image was measured. For intensity of GFAP, Iba1 and CS-56, data were expressed as the intensity ratio of either ipsilateral ROI to the control contralateral ROI. For nuc/cyto YAP intensity, the nuclear boundaries were defined by using the corresponding DAPI stained image, the total nuclear intensity of YAP was divided by the total intensity of YAP outside the nuclear area to obtain the nuc/cyto YAP ratio. For fibronectin area, ROI images (500×1000 µm) were taken adjacent to the scaffold. Data were expressed as the proportional area of fibronectin positive pixels.

Host cell infiltration was assessed at the rostral, central, and caudal areas (350×350 µm) by quantifying the number of cell nuclei and capillaries filled with DAPI^+^ nuclei. Capillaries that contained only single, isolated DAPI^+^ nuclei were classified as non-cell filled. For angiogenesis, data were expressed as the number of blood vessels within the entire scaffold area, which were manually counted from the images obtained. βIII-tubulin or 5-HT labeled axons that crossed virtual lines perpendicular to the rostro-caudal axis was directly counted manually under a fluorescent microscope. These virtual lines were placed at distances of 100 µm and 500 µm away from the hydrogel edge, as well as in the central region. Subsequently, the axon numbers were normalized to the entire scaffold area using the following equation:

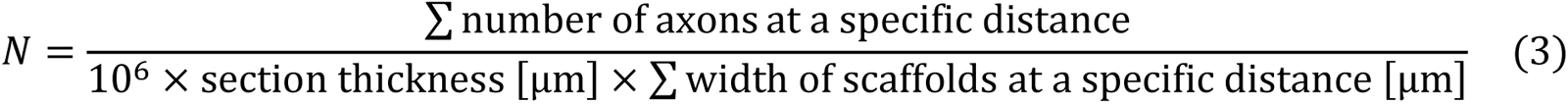

For growth direction of axons, βIII-tubulin labeled images were used. With AngleJ, a plugin for ImageJ, morphological thinning was performed to produce a skeletonized version of the bitmap.[36] Subsequently, the angles were measured relative to a rostral-caudal axis, with 0° representing horizontally growing axons and −90°/90° representing vertically growing axons. The measured angles were then grouped into 36 bins (5° for each bin) ranging from −90° to 90°. Data were expressed as a percentage of axons in each bin relative to the total number of axons observed and the aggregated bins (± 10°, ± 20°)

### Host-implant slices preparation for AFM measurement

4 weeks after ACH scaffold implantation, rats were euthanized with a lethal dose of a ketamine (62.5 mg/kg), xylazine (3.175 mg/kg), and acepromazine (0.625 mg/kg) mixture, diluted in sterile 0.9% saline via *i.p.* injection. The permanent cessation of the circulation was confirmed by subsequent cardiac perfusion with cold (4°C) slicing artificial cerebrospinal fluid (aCSF). The spinal cord tissues integrated with ACH scaffolds were dissected out and kept in cold slicing aCSF for 5min. The composition of slicing aCSF was: 191 mM sucrose, 0.75 mM K-gluconate, 1.25 mM KH2PO4, 26 mM NaHCO3, 4 mM MgSO4, 1 mM CaCl2, 20 mM glucose, 2 mM kynurenic acid, 1 mM (þ)-sodium L-ascorbate, 5 mM ethyl pyruvate, 3 mM myo-inositol, and 2 mM NaOH.[43] Subsequently, a 2 cm long piece of spinal cord integrated with an ACH scaffold was freed from the overlying dura and embedded in 2% low melting point agarose in 0.1 M PB. Then a small agarose block containing the host-implant combination was then glued onto a vibratome platform and 1 cm thick horizontal host-implant slices were cut in cold slicing aCSF with a frequency of 100 Hz and a forward speed of 40 mm/s. The slices were then collected and placed in 35 mm petri dishes treated with BD Cell-Tak (Cell and Tissue Adhesive, BD Biosciences). The petri dishes were then perfused with fresh measuring aCSF (121 mM NaCl, 3 mM KCl, 1.25 mM NaH2PO4, 25 mM NaHCO3, 1.1 mM MgCl2, 2.2 mM CaCl2, 15 mM glucose, 1 mM (þ)-sodium L-ascorbate, 5 mM ethyl pyruvate and 3 mM myo-inositol).[43] The dishes were slowly brought to 37°C for 15–30 min before AFM measurements commenced. This process was conducted less than 1h after sacrificing.

### AFM measurements on host-implant slices

AFM micro-indentation measurements were set up similarly as before. Differently, the indentation entity employed a tipless silicon cantilever (Arrow-TL1, NanoWorld, Neuchatel, Switzerland, spring constants 0.03 Nm^-1^), which was modified by gluing 46.3 µm diameter polystyrene beads (PS/Q-R-B1020, microParticles GmbH, Germany) to the tip of the cantilever via a two-component curing glue (plus schnellfest, UHU, Germany) (Figure S7, Supporting Information). The measurements were performed at 37 °C in measuring aCSF solution. Force–distance curves were taken with an approach speed of 5-10 µm/s produced average indentation depths *δ* 5-15 µm and a set force of 20-30 nN maximally. The boundaries of the host-implant interface were determined by eye. ROIs were selected from the uninjured spinal cord (500×1000 µm, 3 mm away from the scaffold), injured spinal cord (500×500 µm, scaffold adjacent rostrally and caudally), and the center (500×1000 µm) of the implanted scaffold. The apparent Young’s modulus *E* was calculated based equation 1. All curves in each ROI were all pooled together to represent the ROI for statistical analysis.

### Statistical analysis

All data are presented as mean ± standard error of the mean (SEM). Statistical analysis was performed using GraphPad Prism 9. Schematic diagrams in the figures were created with biorender.com. For comparison between two groups, an unpaired Student’s t-test or a Mann-Whitney test (for non-normally distributed data) was employed. For comparisons between multiple groups, a one-way ANOVA followed by a Tukey’s post hoc test, or a Kruskal-Wallis followed by a Dunn’s test (for non-normally distributed data) was utilized. For comparisons with two variables between multiple groups, a two-way ANOVA followed by a Tukey’s post hoc test was performed. A significance criterion of *P* < 0.05 was used.

## Supporting information

Zheng et al Supporting Info

## Acknowledgements

This research was funded by (1) Deutsche Forschungsgemeinschaft (PU 425/4-3, MU 2787/3-3 and WE 2165/4-3) awarded to R.P., R.M. and N.W., (2) Olympia Morata Program Fellowship of the University of Heidelberg Faculty of Medicine awarded to R.P., and (3) Y.Z. sponsored by China Scholarship Council (CSC). We would like to thank Dr. Torsten Müller (JPK Instruments) for technic support on AFM measurements and Dr. Ke Tian for AFM data analysis.

