## Supplementary material for "Biomaterial Scaffold Stiffness Influences the Foreign Body Reaction, Tissue Stiffness, Angiogenesis and Neuroregeneration in Spinal Cord Injury": Zheng et al Supporting Info

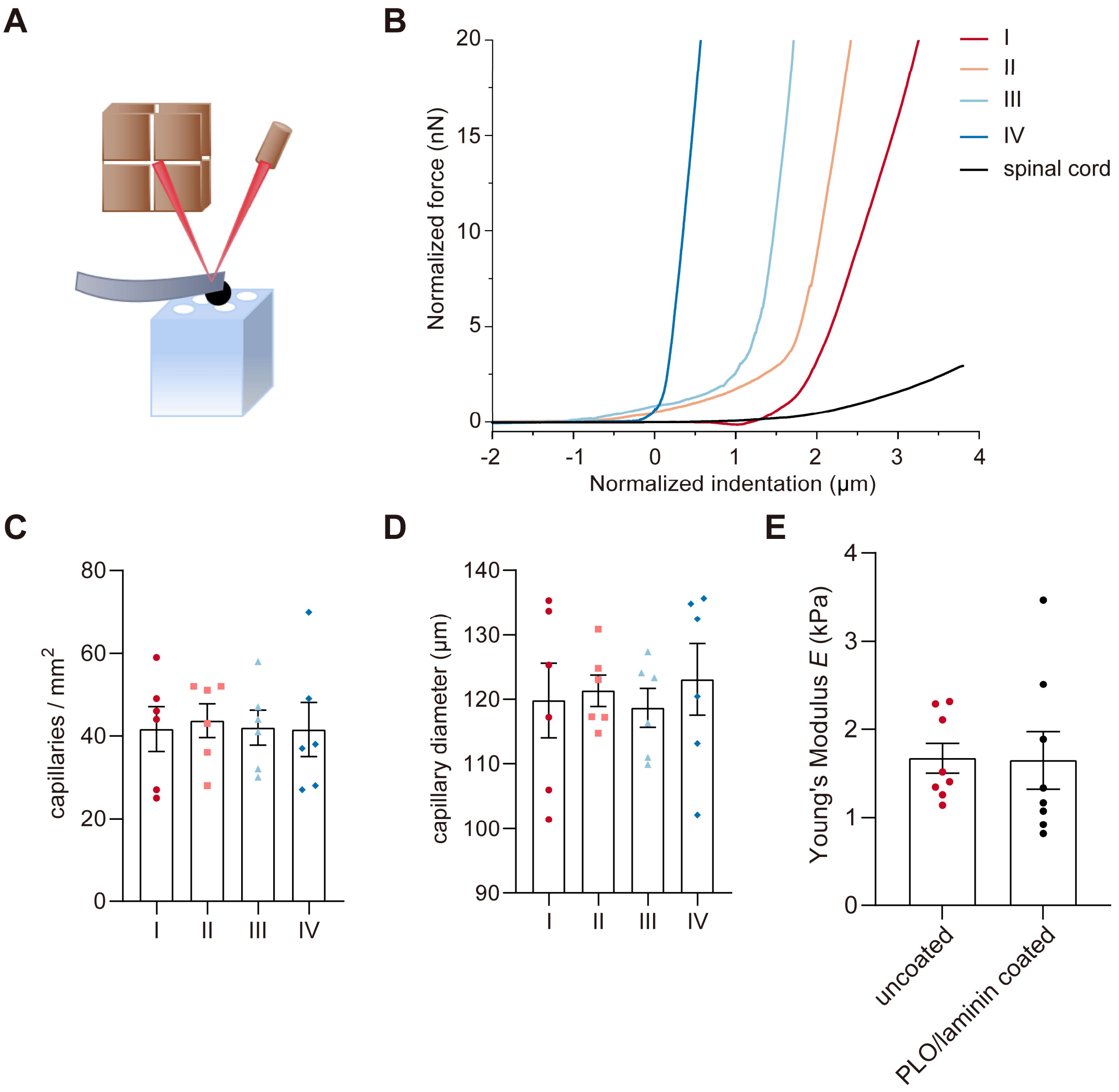


**Figure S1**: Characteristics of stiffness-varied implantable ACH scaffolds. (A) A schematic diagram illustrating the process of AFM-based micro-indentation measurements on implantable ACH cuboids. (B) Representative force-distance curves for each type of scaffold and the spinal cord. (C, D) Structural analysis performed by a light microscope revealed that all types of ACH cuboids exhibited consistent capillary densities of roughly 40 capillaries/mm$^2$ and capillary size of 120 μm (N=7). (E) Quantification of apparent Young's modulus revealed no significant changes in type I ACH cuboids following the PLO/laminin coating process (N=8). Statistical analysis was performed using a one-way ANOVA followed by a Tukey's post hoc test for *C*, *D* and an unpaired t-test for *E* (error bars represent the standard error of the mean).

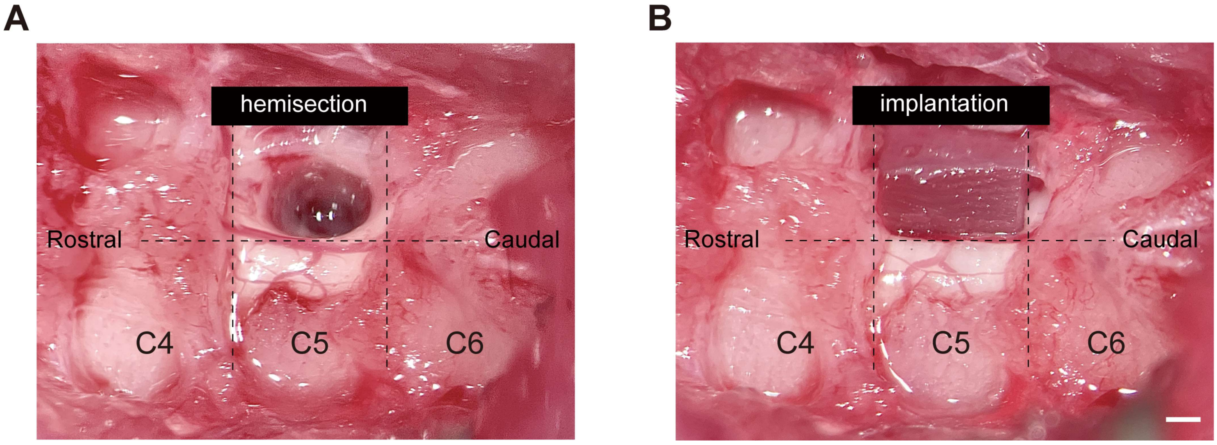


**Figure S2**: *In vivo* surgical procedure. (A) Following a cervical level 5 (C5) lateral hemisection (2×1.3 mm) in the rat spinal cord, (B) the ACH scaffolds (2×1.3×2 mm) were immediately implanted into the lesion site, effectively spanning the gap with the capillaries aligned parallel to the rostral-caudal axis. Scale bar: 500 µm.

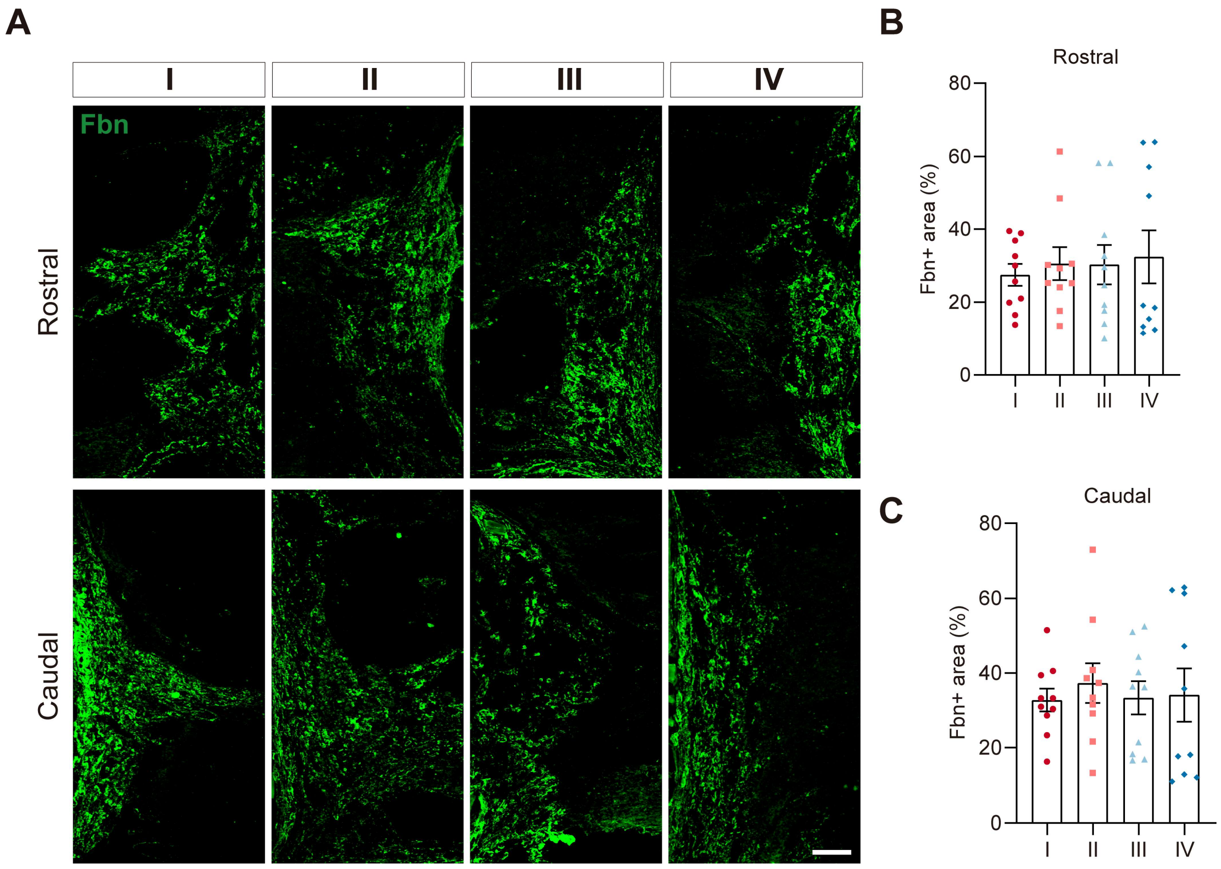


**Figure S3**: Deposition of fibronectin surrounding implanted ACH scaffolds. (A) Immunolabeling of fibronection (Fbn) reveals deposition surrounding each type of implanted scaffold at both rostral and caudal ends. Scale bar: 100 µm. (B, C) Quantification of the proportional area on the rostral or caudal end of the scaffolds for each group (N=10). Statistical analysis was conducted using a Kruskal-Wallis test followed by a Dunn's test (error bars: standard error of the mean).

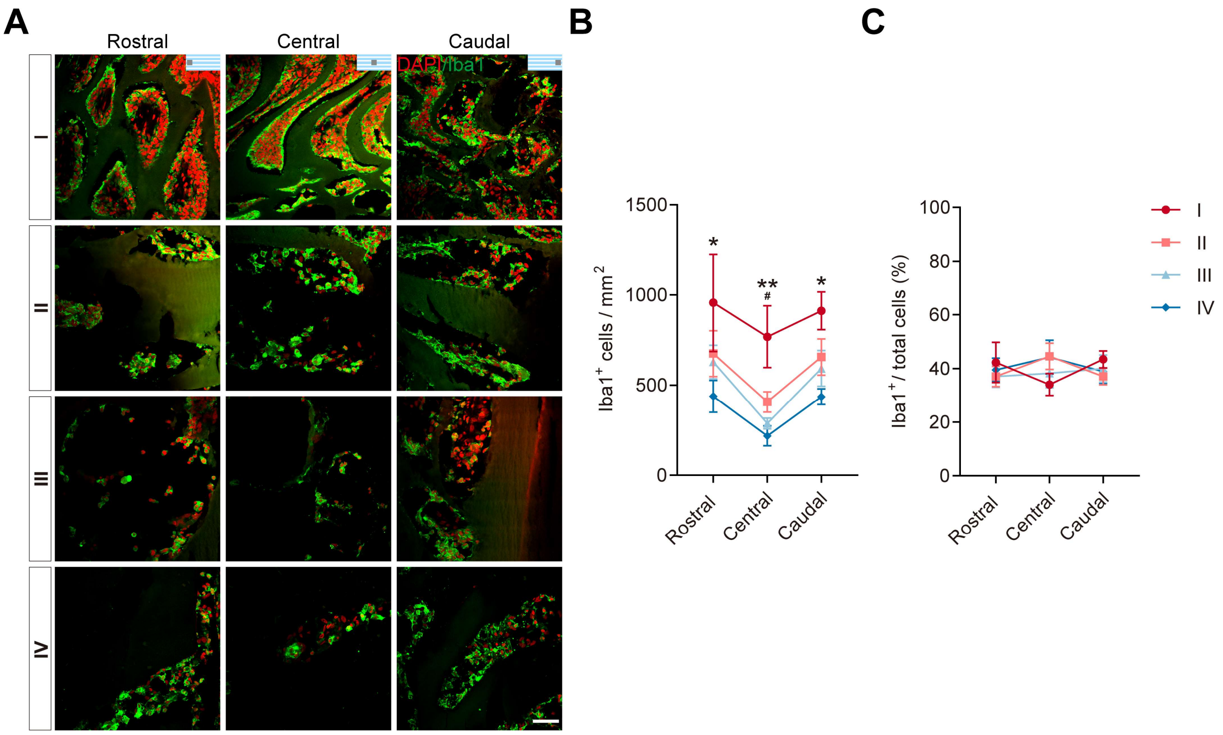


**Figure S4**: Infiltration of microglia/macrophages within scaffold capillaries. (A) Co-immunolabeling of DAPI and Iba1 reveals microglia/macrophage infiltration into capillaries in each group of implanted scaffolds, with the three columns corresponding to the rostral, central, and caudal aspects of the scaffolds, as illustrated by the schematic in the upper right corner in the first row. Scale bar: 50 μm. (B, C) Quantifying the number of infiltrated microglia/macrophages per square millimeter and the percentage of microglia/macrophages in each aspect of the scaffold for each group (N=10). Statistical analysis was conducted using a two-way ANOVA followed by a Tukey's post hoc test (* $P_{I\ vs\ IV} < 0.05$, ** $P_{I\ vs\ IV} < 0.01$, # $P_{I\ vs\ III} < 0.05$, error bars: standard error of the mean).

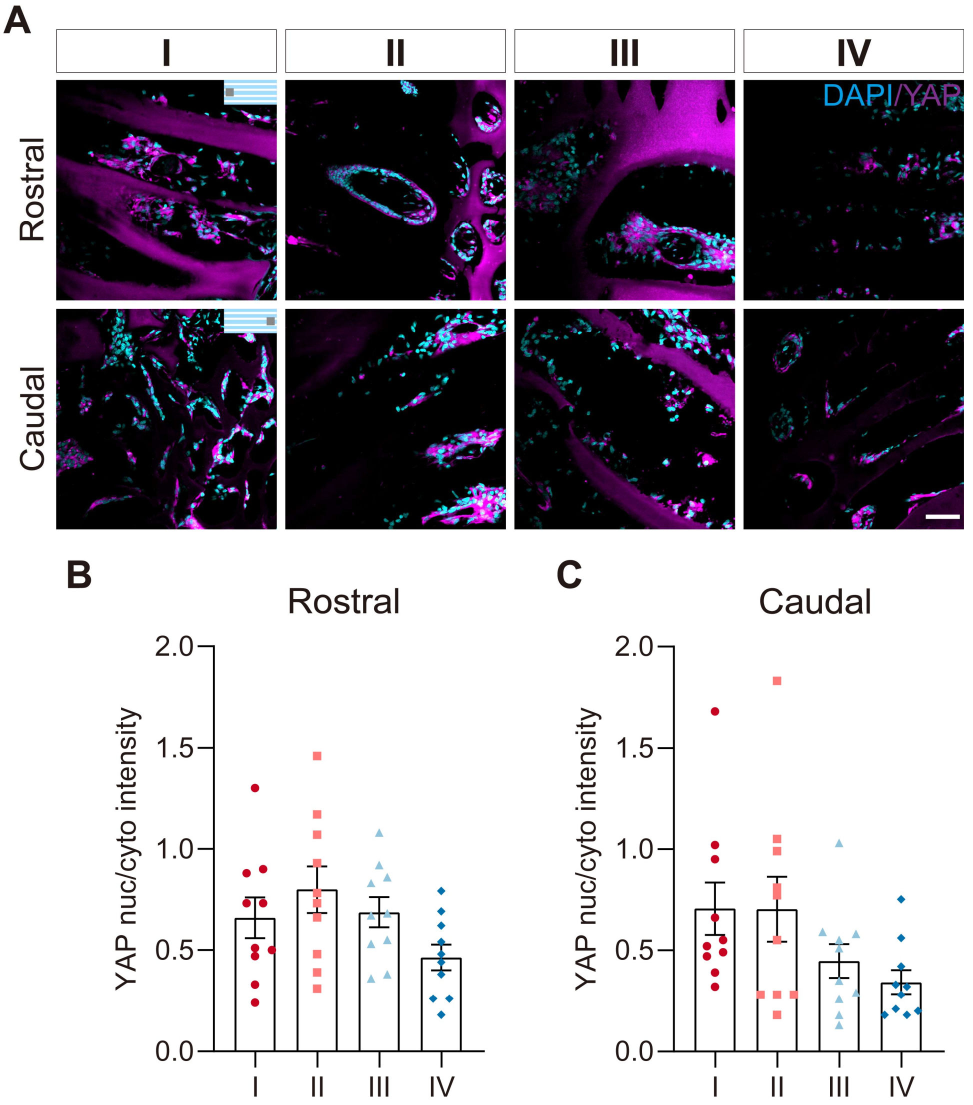


**Figure S5**: YAP nuclear translocation within the entries of the ACH scaffolds does not differ. (A) Co-immunolabeling of YAP and DAPI in the rostral and caudal aspect of scaffold capillaries for each group. Scale bar: 50 µm. (B, C) Quantifying nuc/cyto YAP intensity within the rostral and caudal capillaries of the scaffold (N=10). Statistical analysis was conducted using a one-way ANOVA followed by a Tukey's post hoc test (error bars: standard error of the mean).

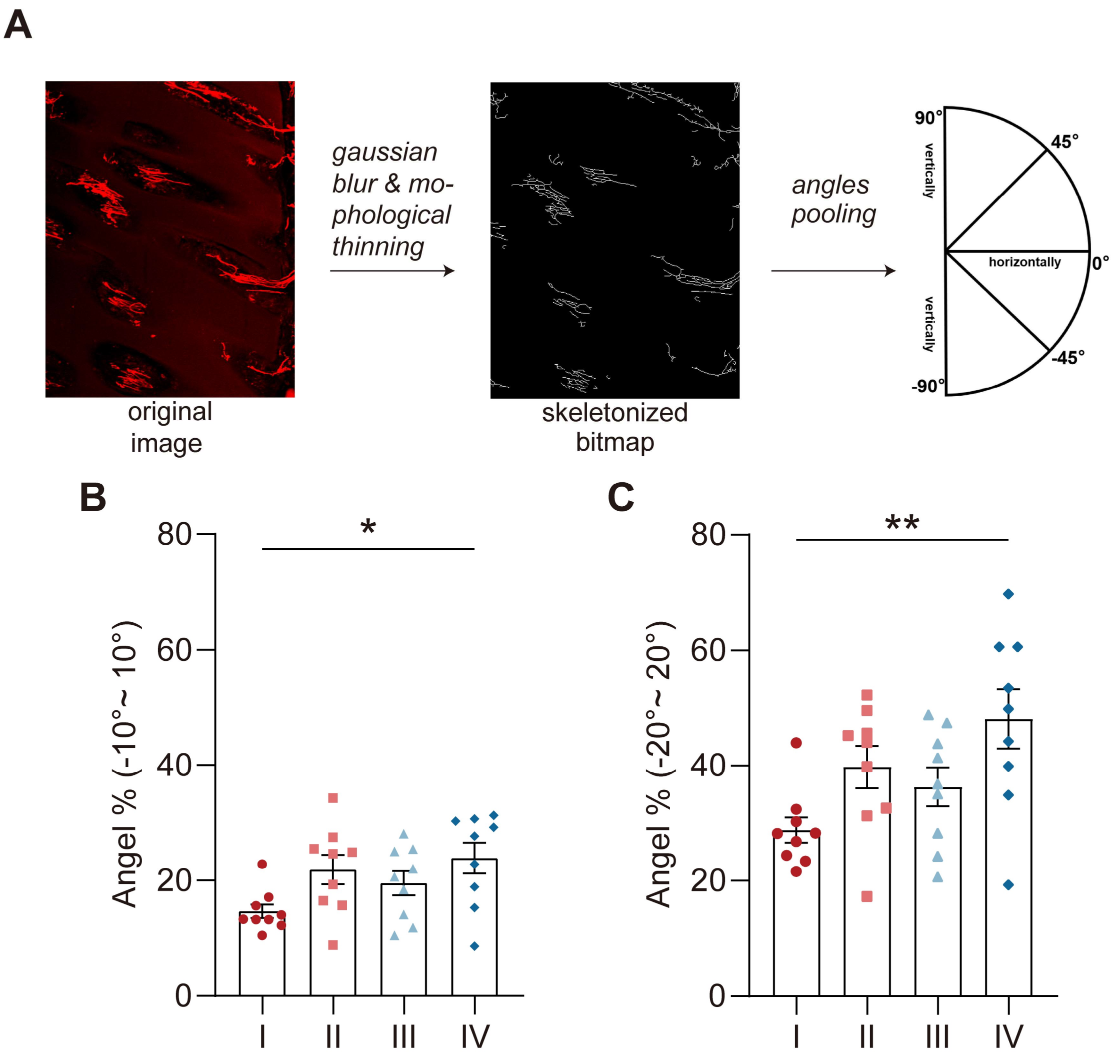


**Figure S6**: Orientation measurement of regrowing axons within implanted scaffolds. (A) For the determination of axonal growth direction, βIII-tubulin labeled images were utilized. Using the AngleJ plugin for ImageJ, morphological thinning was performed to create a skeletonized version of the bitmap. The measured angles were then grouped into 36 bins (5° for each bin) spanning from -90° to 90°. (B, C) Quantification of rostral-caudal orientations (±10° or ±20°) of regrowing axons within each type of scaffold (N=10). Statistical analysis was conducted using a one-way ANOVA followed by a Tukey's post hoc test (*$P < 0.05$, **$P < 0.01$, error bars: standard error of the mean).

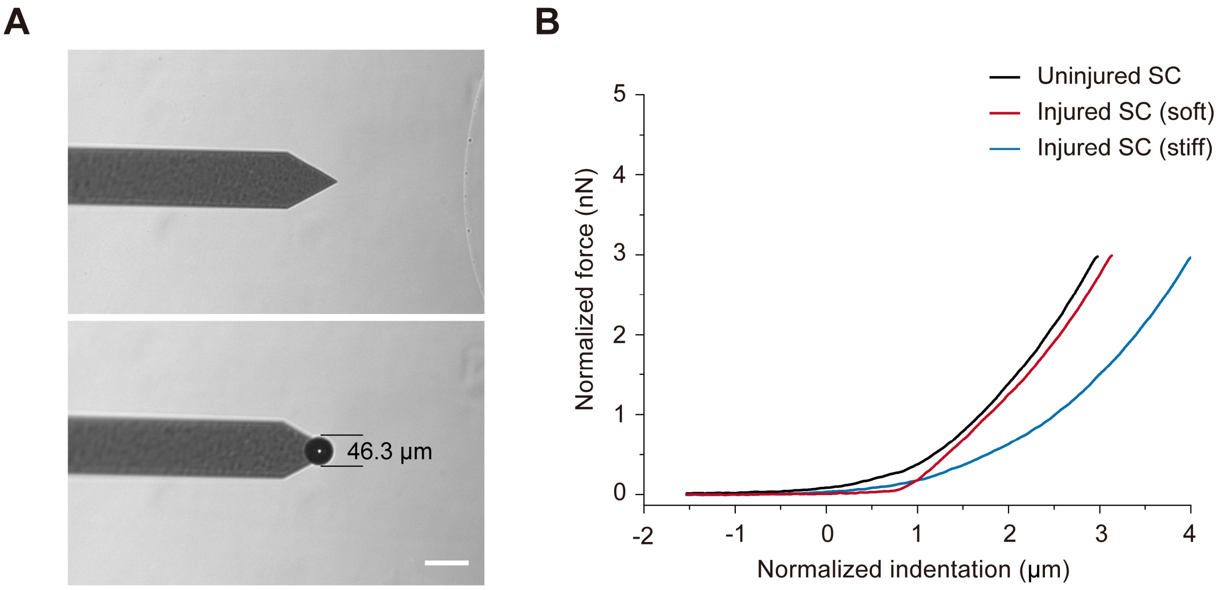


**Figure S7** AFM measurements of the host-implant slices 4 weeks post-implantation. (A) For determining the mechanics of the very soft spinal cord tissue, a large sphere tip with a diameter of around 46.3 µm was used to glue on the very top of the tipless cantilever. Scale bar: 100 µm (B) Representative force-distance curves for R1 vs R2/R2' on host-implant slices.

**Table S1** Antibody list for immunohistochemistry.

| Antibody | Host | Dilution | Company | Target |
|---|---|---|---|---|
| anti-GFAP | rabbit | 1:1000 | Dako | Astrocytes |
| anti-GFAP | guinea pig | 1:1000 | PROGEN | Astrocytes |
| anti-Iba1 | rabbit | 1:500 | Wako | Microglia/Macrophages |
| anti-C3 | rabbit | 1:300 | Abcam | C3 components |
| anti-CS-56 | mouse | 1:200 | Sigma-Aldrich | CSPGs |
| anti-YAP | mouse | 1:200 | Santa Cruz | YAP |
| anti-Fibronectin | rabbit | 1:100 | Abcam | Fibronectin |
| anti-CD31 | goat | 1:300 | R&D Systems | Angiogenesis |
| anti-βIII-tubulin | mouse | 1:1000 | Promega | Axons |
| anti-5-HT | rabbit | 1:2000 | Immunostar | Serotonergic axons |
| DAPI | - | 1:2000 | Sigma-Aldrich | Cell Nuclei |
| Alexa Fluor 488 | Donkey | 1:300 | Invitrogen | Anti-mouse |
| Alexa Fluor 488 | Donkey | 1:300 | Invitrogen | Anti-rabbit |
| Alexa Fluor 594 | Donkey | 1:300 | Invitrogen | Anti-mouse |
| Alexa Fluor 594 | Donkey | 1:300 | Invitrogen | Anti-rabbit |
| Alexa Fluor 594 | Donkey | 1:300 | Invitrogen | Anti-goat |
| Cy5 | Donkey | 1:500 | Jackson | Anti-rabbit |
| Cy5 | Donkey | 1:500 | Jackson | Anti-guinea pig |

**Table S2** Quantification of the contralateral side for GFAP and Iba1 immunostaining.

| Group | Mean gray value (GFAP) | Mean gray value (Iba1) |
|---|---|---|
| I | 19.85 | 32.80 |
| II | 19.39 | 28.32 |
| III | 20.62 | 27.67 |
| IV | 17.46 | 31.89 |
| Sample size | N = 10 | N = 10 |
| One way-ANOVA | P = 0.8375 | P = 0.4781 |
